# A fat-tissue sensor couples growth to oxygen availability by remotely controlling insulin secretion

**DOI:** 10.1101/348334

**Authors:** Michael J. Texada, Anne F. Joergensen, Christian F. Christensen, Daniel K. Smith, Dylan F.M. Marple, E. Thomas Danielsen, Sine K. Petersen, Jakob L. Hansen, Kenneth A. Halberg, Kim F. Rewitz

## Abstract

Organisms adapt their metabolism and growth to the availability of nutrients and oxygen, which are essential for normal development. This requires the ability to sense these environmental factors and respond by regulation of growth-controlling signals, yet the mechanisms by which this adaptation occurs are not fully understood. To identify novel growth-regulatory mechanisms, we conducted a global RNAi-based screen in *Drosophila* for size differences and identified 89 positive and negative regulators of growth. Among the strongest hits was the *FGFR* homologue *breathless* necessary for proper development of the tracheal airway system. *Breathless* deficiency results in tissue hypoxia (low oxygen), sensed primarily in this context by the fat tissue. The fat, in response, relays this information through release of one or more humoral factors that remotely inhibit insulin secretion from the brain, thereby restricting systemic growth. Thus our findings show that the fat tissue acts as an oxygen sensor that allows the organism to reduce its growth in adaptation to limited oxygen conditions.

---

Multicellular organisms must adapt their metabolism and growth to meet the limitations imposed by their environment. These processes are regulated by internal genetic and physiological pathways, such as growth-factor or steroid-signaling pathways, and these in turn are influenced by internal and external cues. Among these are nutrient availability, which promotes growth through the action of the insulin/insulin-like growth factor (IGF) pathways^1–3^, and oxygen sufficiency, which is also essential for the growth and development of animals^4, 5^. The signaling pathways that regulate these fundamental aspects of development are evolutionarily ancient and are well-conserved.

Signaling through Insulin and IGF is the major systemic nutrient-dependent growth regulator and acts through conserved receptors and downstream pathways to promote cellular growth^3, 6, 7^. Many connections between nutritional cues and systemic insulinergic regulation have been elucidated in the fruit fly *Drosophila melanogaster*. In this model organism, several *Drosophila* insulin-like peptides (DILPs or simply “insulin” below), primarily DILP2, −3, and −5, are released into circulation from the insulin-producing cells (IPCs) of the brain^8–10^. Despite their different location, the IPCs are functionally homologous with the insulin-producing pancreatic β-cells of mammals^9^. These circulating DILPs all signal through a single insulin receptor (InR) to regulate both metabolism and growth. The expression and release of individual DILPs are regulated in part by nutritional information relayed through the fat body, a major organ for nutrient storage and metabolism that also has endocrine functions and which is analogous to vertebrate adipose and liver tissues. The fat tissue secretes insulinotropic (the cytokine Unpaired-2, a functional analog of the mammalian adipose-derived hormone Leptin^11^; the peptide hormones CCHamide-2^12^, FIT^13^, and Growth-Blocking Peptides^13, 14^; and the protein Stunted^15^) and insulinostatic (the TNF-α homolog Eiger^16^) factors, many of these in response to nutrient-dependent activity of the Target of Rapamycin (TOR) pathway. Thus, this tissue is a central nexus for nutritional signals that mediate inter-organ communication between the fat and the brain, which allows organisms to reduce their growth rate in adaptation to nutritional deprivation.

Steroid hormones make up a second class of growth-regulatory signals that also control developmental transitions including maturation, the transition to adulthood, which terminates the juvenile growth period in many animals. In *Drosophila*, this developmental transition is regulated by the steroid hormone ecdysone, which initiates metamorphosis into the adult. Ecdysone is produced and released by the prothoracic gland (PG) in response to the neuropeptide prothoracicotropic hormone (PTTH) and insulin^17–19^. During development, basal levels of ecdysone suppress systemic body growth, while high-level ecdysone peaks limit the larval growth period by triggering pupariation.

In addition to nutrients, organisms require oxygen for growth and development. Organisms have therefore developed oxygen-sensing mechanisms that allow them to adapt their metabolism and growth to low oxygen concentrations. In many organisms, hypoxia (low oxygen level) slows growth, leading to delayed development and smaller adults. In *Drosophila* and other insects, hypoxia restricts growth and whole-body size^4, 20–24^. Furthermore, in humans, the limited oxygen associated with high-altitude living has been linked to slowed growth and developmental delay^25, 26^. This effect arises at oxygen concentrations above those that compromise the basic metabolic rate^27, 28^, suggesting that this is not simply an effect of an insufficiency of O_2_ to support aerobic respiration, but rather an active adaptation under genetic control. The conserved transcription factor Hypoxia-Inducible Factor 1 alpha (HIF-1a) is the key regulator required for adaptive hypoxia responses^29^. At a rate proportional to oxygen levels, the enzyme HIF-1a Prolyl Hydroxylase (Hph) hydroxylates specific residues in the oxygen-dependent degradation (ODD) domain of HIF-1a, leading to its degradation. Under hypoxic conditions, oxygen is insufficiently available for this process, and HIF-1a becomes stable and dimerizes with the constitutively expressed HIF-1 beta (HIF-1b), translocates to the nucleus, and induces target-gene expression. While it is well established that nutrients mainly affect growth through insulin, the precise mechanism by which limited oxygen restricts systemic growth as an adaptation for survival under low oxygen levels remains to be determined.

To identify signaling mechanisms with roles in regulating systemic growth and body size, we conducted an RNAi-based screen of 1,845 genes encoding potential secreted factors and their receptors. The receptor tyrosine kinase gene *breathless (btl)*^30, 31^, which encodes an FGF-receptor (FGFR) homolog, was one of the strongest hits. This receptor is expressed by cells of the tracheae, the gas-delivery tubules of the insect respiratory system. Its FGF-like ligand Branchless (Bnl) is produced by target tissues in a stereotyped pattern during development^32^ and in response to hypoxia during larval growth^33, 34^ to guide the tracheal terminal branches toward target tissues. When this system is perturbed, internal tissues experience hypoxia. We found that tissue hypoxia – actual hypoxia induced by manipulation of Btl-mediated tracheal growth or external O_2_ levels, or genetically induced fictive hypoxia adaptation via Hph or HIF-1a perturbations – altered the expression of *Dilp2*, −*3*, and −*5* in the IPCs and blocked DILP release, leading to reduced systemic insulin signaling. We further report that the primary sensor of internal oxygen levels is the fat body, which in response to hypoxia releases one or more as-yet unidentified insulinostatic humoral factors that induces these changes in IPC physiology. Understanding the changes brought about by hypoxia in this model system may allow greater understanding of human disease associated with tissue-hypoxia such as obesity and diabetes.

## Results

### *In-vivo* RNAi screen identifies signals affecting body size

We undertook a genetic RNAi-knockdown^35, 36^ screen targeting 1,845 genes encoding the *Drosophila* secretome and receptome (Fig. 1A, Supplemental Table 1). Ubiquitous gene knockdown using the *daughterless-GAL4* driver (*da*>) identified 89 genes whose manipulation resulted in significantly large or small pupae (Fig. 1B and 1C). Among these hits were several genes already known to regulate body or tissue growth (Fig. 1C and 1D), including those encoding the insulin receptor (InR)^37^, Anaplastic lymphoma kinase (Alk)^38^ and its ligand Jelly Belly (Jeb)^39^, the epidermal growth-factor receptor (EGFR)^40^, and the PTTH receptor Torso^41^, thus validating our approach. Genes important for ecdysone signaling (*knirps*, *Eip75B*, *Npc2g*, and *Torso*^41–44^) were among the hits that increased pupal body size, consistent with the growth-limiting effects of ecdysone. On the other hand, knockdown of the growth-promoting *InR* produced a strong decrease in body size. Among the strongest hits associated with reduced growth, we also identified FGF-pathway signaling (Fig. S1). The main hit, the *FGFR* ortholog *breathless* (*btl*), and its FGF-ligand Branchless (Bnl), are critical for proper tracheation of tissues to allow for gas exchange and oxygen delivery^30, 31^.

### The FGF receptor Breathless affects systemic growth through effects on tracheae

Ubiquitous *btl* knockdown (*da>btl-RNAi*) caused a ~30% reduction in pupal size (Fig. 2A and 2B). Likewise, whole-body knockdown of the ligand *bnl* strongly reduced growth. Knockdown of the upstream transcription factor *trachealess* (*trh*)^45^, which is also required for tracheal formation, reduced body size to a lesser degree, perhaps because of low RNAi efficiency combined with the activity of Trh-independent feedback signaling later in tracheal development^46^. The main *btl*-expressing tissue is the tracheal system, but the gene is also active in other cell types such as glia^30^. To identify the tissue causing the size phenotype, we assayed the effect of *btl* RNAi in the tracheae (*btl>btl-RNAi*) as well as in the nervous system (*elav>*), muscles (*Mef2>*), and fat body (*ppl>*). Only tracheal knockdown of *btl* led to reduced body size (Fig. 2C); the specificity of the RNAi effect was confirmed with two additional independent RNAi lines against *btl* that produced similar growth phenotypes (Fig. S2A). Knockdown of *btl* in the tracheae delayed pupariation, prolonging the larval growth period, suggesting that the body-size reduction was the result of a reduced growth rate (Fig. S2B). To investigate this, we determined the growth rate during the third larval instar (L3) and found that loss of *btl* in the trachea slows the growth of the entire animal (Fig. 2D). Together these findings show that loss of *btl* in the tracheal system leads to systemic growth reduction, possibly as a consequence of oxygen restriction.

**Fig. 1.**
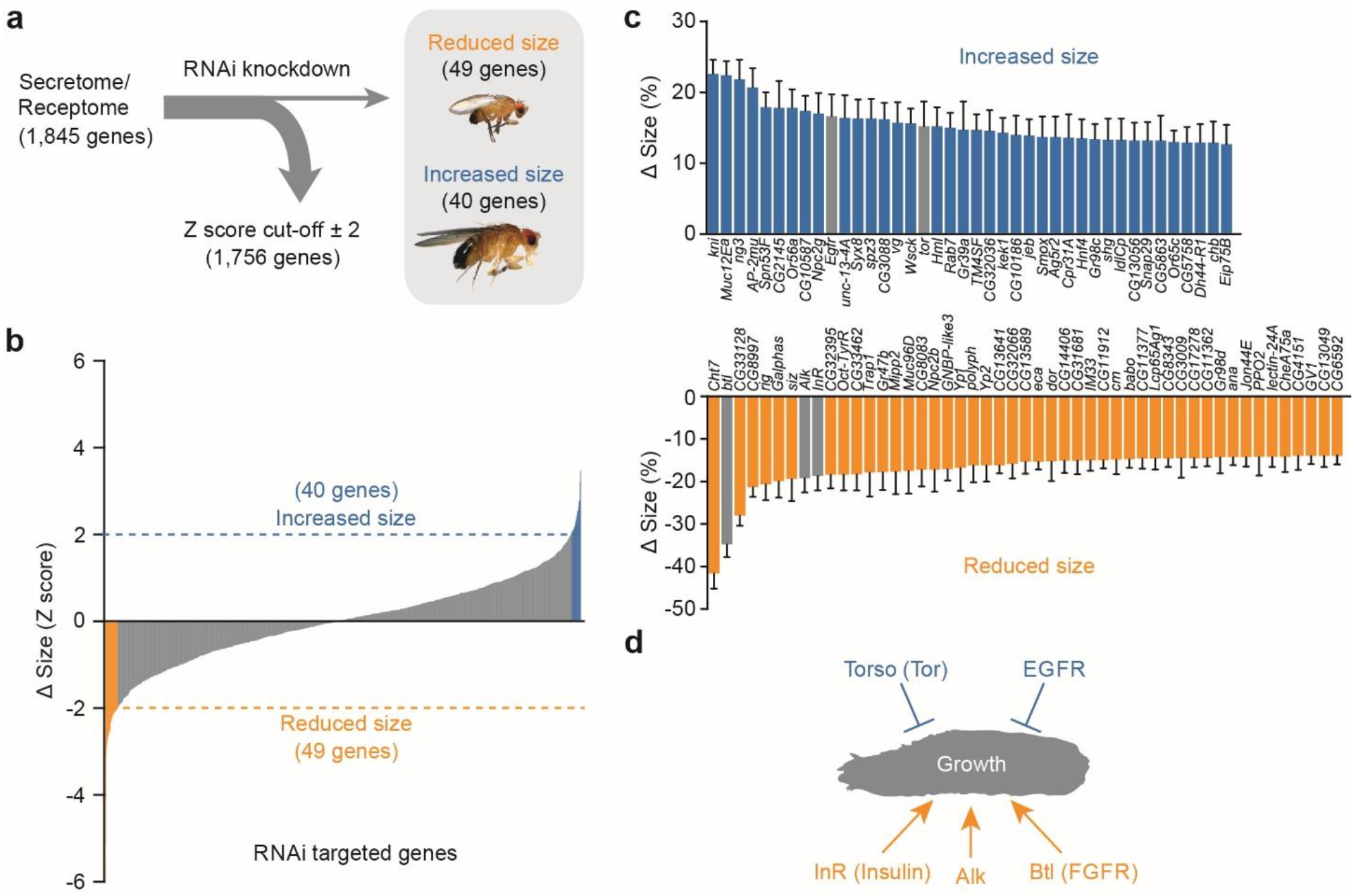
*In-vivo* RNAi screen of the secretome and receptome for factors regulating body size. **a** General scheme of the RNAi screen of 1,845 genes of the *Drosophila* secretome and receptome selected by gene-ontology analysis. UAS-inducible RNAi constructs against these genes were expressed ubiquitously using *da-GAL4* for global knockdown, and pupal size was determined. **b-c** Distribution of the pupal sizes from the screen revealed (**b**) 89 gene hits that significantly resulting size increase (**c**: blue, top) or decrease (**c**: orange, bottom) body size (a *Z*-score > ± 2 was used as a cut off) by the comparison to the mean of all lines. **d** Genes and pathways with known size-regulatory function identified in the screen, as well as *breathless* (*btl*), are marked with arrows.

**Fig. 2.**
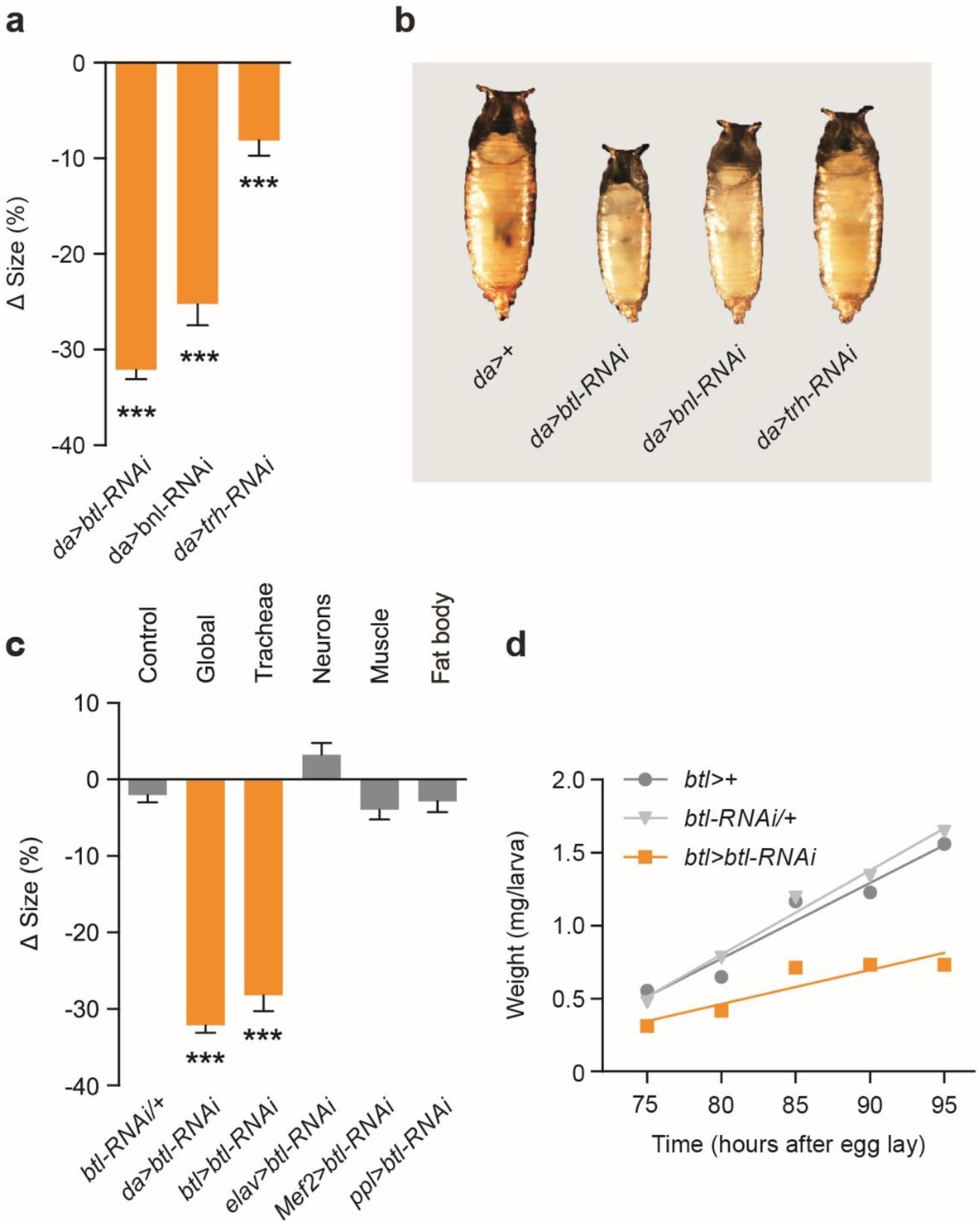
Loss of the FGF receptor *breathless* in the tracheae reduces growth and body size. **a-b** Quantification of pupal sizes (**a**) and representative images (**b**) of animals with ubiquitous knockdown of *breathless* (*btl)*-pathway components compared to the controls (*da>*+; *da>* crossed to wild type, *w^1118^*). Values are percent change in pupal size versus control. **c** Knockdown of *btl* in whole animals (*da>btl-RNAi*) or tracheae alone (*btl>btl-RNAi*), but not in neurons (*elav>btl-RNAi*), muscle (*Mef2>btl-RNAi*), or fat-body tissue (*ppl>btl-RNAi*), leads to small animals compared to *da>+* controls. Shown also is the RNAi control (*btl-RNAi*/+; *btl-RNAi* crossed to wild type, *w^1118^*). Values are percent change in pupal size versus the *btl>*+ controls (*btl>* crossed to wild type, *w^1118^*). **d** Trachea-specific knockdown of *btl* reduces larval growth rates compared to driver (*btl>*+) and RNAi controls (*btl-RNAi/+*). Statistics: one-way ANOVA with Dunnett’s multiple-comparisons test. ***, P<0.001, compared to the control.

### The *breathless*-RNAi size phenotype is mediated by insulin signaling

Among the primary regulators of systemic growth in *Drosophila* are the DILPs^10, 47^. To assess whether the body-size phenotype of *btl* knockdown was mediated by alterations in insulin signaling, we first examined the effects of reducing insulin signaling on development and body size. We found that global reduction of insulin signaling by *InR* knockdown or inhibition of the TOR-dependent fat-body nutrient sensor by tissue-specific expression of the TOR inhibitors *TSC1* and *TSC2* (*TSC1/2*), both strongly reduced pupal size (Fig. 3A) and prolonged development (Fig. S3A), phenocopying the trachea-specific loss of *btl*. To identify any alterations in insulin expression caused by *btl>btl-RNAi*, we determined the level of *Dilp2*, −*3*, and −*5* gene expression in feeding third-instar larvae, ~24 hours prior to pupariation. Indeed, a very strong reduction (>90%) in *Dilp3* expression was observed in RNAi animals, accompanied by a smaller decrease in *Dilp5* transcripts (Fig. 3B).

Since insulin activity is also regulated at the level of DILP release^48^, we investigated whether tracheal *btl* loss induced IPC retention of DILP2, −3, and −5. Animals with trachea-specific *btl* knockdown exhibited an accumulation of these DILPs in their IPCs (Fig. 3C), despite the reduced expression of *Dilp3* and *Dilp5*. This indicates a strong decrease in DILP secretion in larvae with reduced expression of *btl* in the trachea. Presumably, this retention of insulin would reduce signaling through InR in peripheral tissues. To assess this, we measured downstream insulin-signaling mediators. Insulin signaling leads to the phosphorylation of the kinase Akt (pAkt) and suppression of *InR* as part of a feedback circuit regulating cellular sensitivity to insulin^49, 50^. We observed *InR* upregulation and pAkt reduction in *btl>btl-RNAi* animals (Fig. 3D, 3E, and 3F), indicating a reduction in systemic insulin signaling, presumably arising from blocked IPC DILP release. To determine whether the observed reduction in insulin signaling was the direct cause of the observed size phenotype, we asked whether rescuing circulating insulin levels by overexpression of *Dilp2* would also rescue growth. Ubiquitous expression of *Dilp2* using *da-GAL4* caused lethality, so we used the weaker ubiquitous *armadillo-GAL4* driver *(arm>)* and found again that *btl* knockdown led to reduced body size (Fig. 3G). Overexpression of *Dilp2* in this context rescued the size defect (Fig. 3G), supporting the hypothesis that the size phenotype is caused by a systemic reduction in insulin signaling downstream of Btl.

An alternative explanation for the observed reduced body size involves increased ecdysone signaling, which antagonizes insulin signaling and suppresses larval body growth^17^. However, under *btl* knockdown we observed reduced expression of the ecdysone-induced gene *E75B*, which indicates reduced ecdysone-induced growth restriction (Fig. S3B). Thus, the reduced body size observed with *btl* knockdown animals is mediated by effects in the tracheal system that lead to altered expression and reduced secretion of DILP2, −3, and −5 from the IPCs, resulting ultimately in reduced insulin-driven systemic growth.

**Fig. 3.**
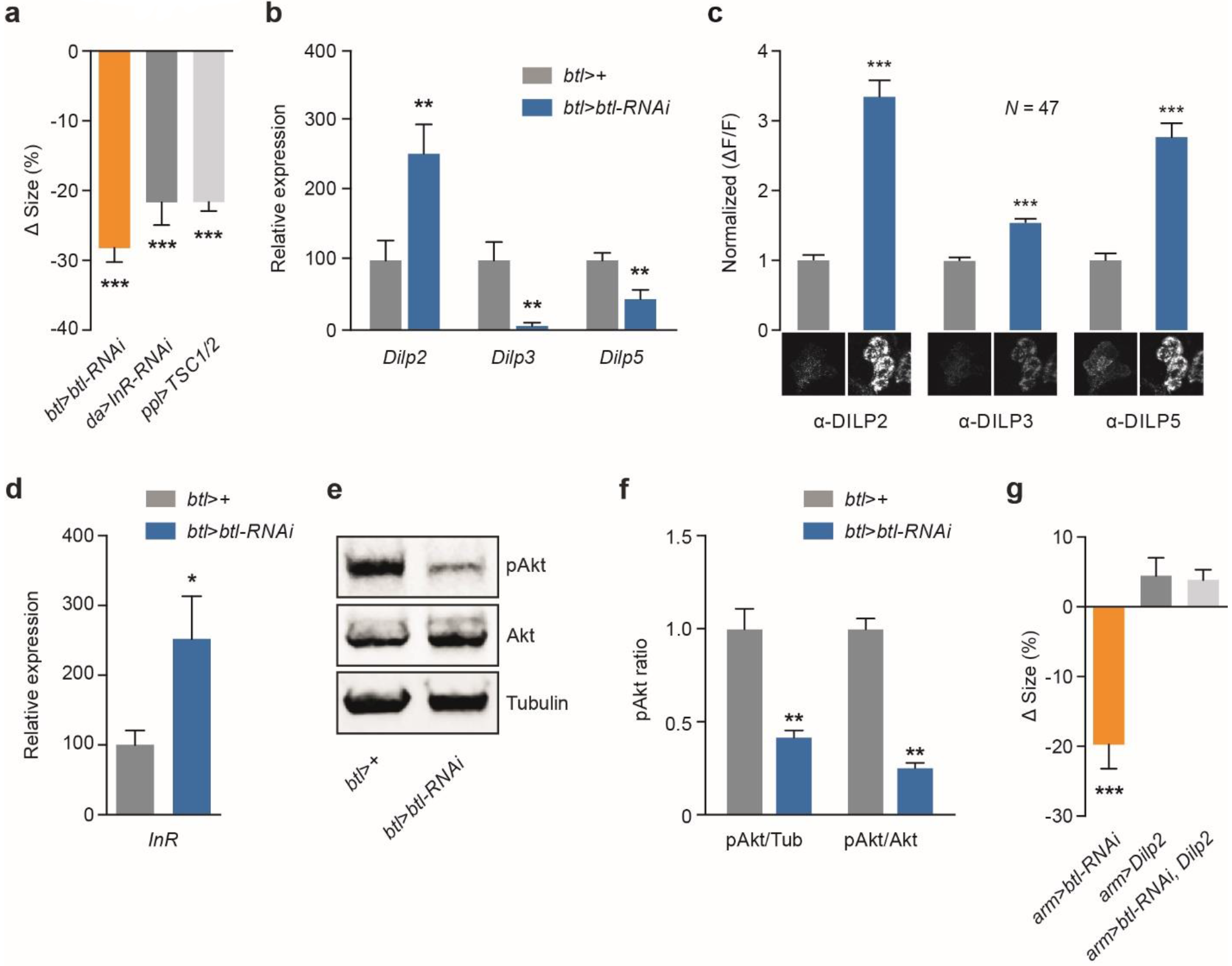
Breathless controls systemic growth through modulation of DILP signaling. **a** Trachea-specific knockdown of *breathless* (*btl)* mimics loss of insulin signaling (ubiquitous driven knockdown of the insulin receptor *InR*; *da>InR-RNAi*) and fat-body inhibition of TOR (*ppl>TSC1/2*), which remotely inhibits release of insulin-like peptides (DILPs) from the insulin-producing cells (IPCs). Values are percent change in pupal size versus the *btl>*+ controls (*btl>* crossed to wild type, *w^1118^*). **b-c** Levels of whole-animal *Dilp* transcripts (**b**) and DILP peptides in the IPCs (**c**) in *btl>btl-RNAi* animals with knockdown of *btl* in the trachea compared to *btl>+* controls. Representative images of DILP2, −3, and −5 immunostainings are shown below. **d** Increased expression of *InR* in *btl>btl-RNAi* animals compared to *btl>+* controls suggests reduced systemic insulin signaling. **e-f** Immunoblotting showing whole-animal levels of phosphorylated Akt (pAkt) kinase (**e**) in *btl>btl-RNAi* animals compared to *btl>+* controls, quantified in (**f**) from three independent replicates times three whole larvae each and normalized to Tubulin (Tub) or Akt levels. Akt and Tubulin were used as loading controls. **g** Ectopic expression of *Dilp2* rescues the *btl*-knockdown size phenotype. Ubiquitous weak knockdown of *btl* using *armadillo-GAL4* (*arm>btl-RNAi*) leads to small pupae (left). Co-expressing *Dilp2* rescues this size phenotype (right), while having no significant effect on control animals (middle). Values are percent change in pupal size versus *arm>+* controls. Statistics: one-way ANOVA with Dunnett’s test for multiple-comparisons and Student’s t-test for pairwise comparisons. *P>0.05, **P>0.01, ***P>0.001, compared to the control.

### *Breathless* knockdown leads to reduced tracheal growth and tissue hypoxia

In the tracheae, Btl acts to increase growth towards tissues experiencing hypoxia, because these tissues the FGF-like ligand Bnl. Thus, knockdown of *btl* would be expected to lead to reduced tracheal branching and oxygen supply to internal tissues. To confirm this, we observed tracheae associated with the fat body and quantified the number of branches and the total length of tracheal tube^51^. The tracheae of control animals display extensive terminal branches, whereas tracheae of *btl>btl-RNAi* animals exhibit significantly fewer of these fine processes (Fig. 4A). Furthermore, the total length of the fat-body tracheal branch was greatly reduced in animals with reduced expression of *btl* in the trachea (Fig. 4B), suggesting reduced oxygen supply leading to tissue hypoxia. To investigate this possibility, we examined the expression of *bnl*, which is upregulated in response to hypoxia to drive tracheal outgrowth towards oxygen-demanding tissues^33^. Loss of *btl* in the trachea led to strong *bnl* upregulation (Fig. 4C), indicating that internal tissues were indeed experiencing hypoxia. We rationalized therefore that these animals would be sensitized to low-oxygen conditions. To test this, we reared control and animals with knockdown of *btl* in the trachea under hypoxic conditions (5% O_2_, *vs*. the normal 21% atmospheric level) and measured their survival to the pupal stage. Most control animals survived this environment, reflecting their ability to adapt to reduced oxygen levels, whereas none of the *btl-RNAi* animals survived to pupariation (Fig. 4D), presumably dying from hypoxia-induced damage. Taken together, our data suggest that the insulinergic changes described above operate downstream of tissue hypoxia to adapt growth to oxygen availability.

To investigate whether the tracheae themselves were under the control of insulin, we assessed the effects of altering insulin signaling in this tissue. Although overexpression of *Dilp2* rescued body growth of *arm*>*btl-RNAi* animals (Fig. 3G), it did not restore the growth of their tracheae (Fig. S3C). Furthermore, tracheal knockdown of *InR* or *Akt* had no effect on systemic growth or pupal size, whereas loss of TOR signaling in the tracheae caused a strong reduction in body size (Fig. S3D). Thus, our findings suggest that insulin works downstream of the growth of the tracheal system and oxygen levels, mediating their effects on systemic growth, and that the growth of tracheae themselves is insulin-independent.

**Fig. 4.**
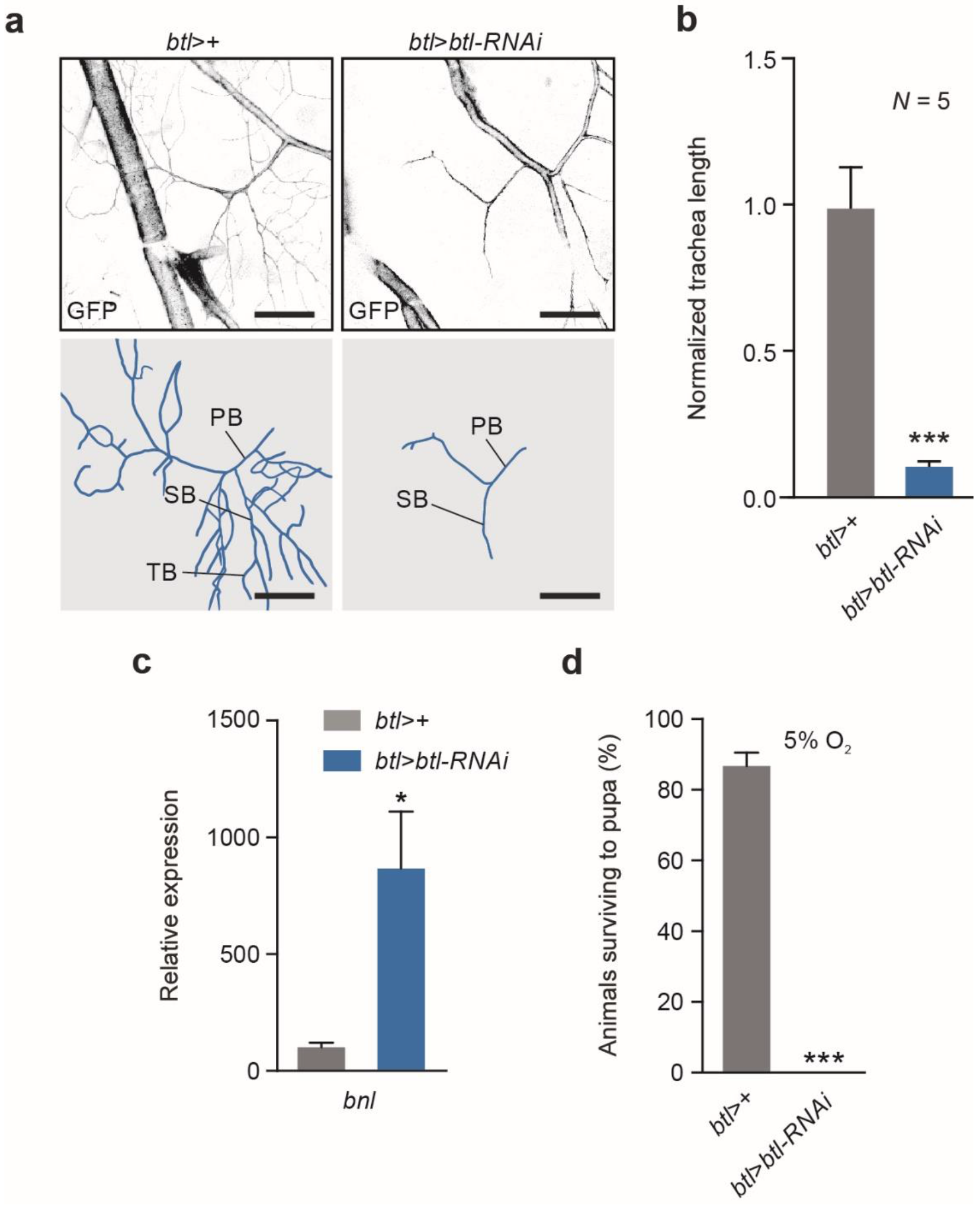
Airway branching is reduced in animals with knockdown of *breathless*. **a-b** Knockdown of *breathless* (*btl)* in tracheae leads to a reduction in branching (right) of the terminal tracheal cells of the fat body branch compared to the *btl>+* control (*btl>* crossed to wild type, *w^1118^*). Representative images (**a**) and quantification (**b**) of the fat-body branch of larval tracheal terminal cells are shown. Scale bar, 50 μm. PB, primary branch; SB, secondary branch; TB, tertiary branch. **c** Whole boyd transcript levels of *branchless* (*bnl*), encoding the hypoxia-inducible FGF ligand of Btl, is strongly upregulated in larvae with trachea-specific *btl* knockdown, suggesting that tissues are experiencing hypoxia in these animals. **d** Loss of *btl* renders animals unable to survive in a low-oxygen environment (5% O_2_ levels), while most *btl>+* controls develop to pupariation. Statistics: Student’s t-test for pairwise comparisons. *P<0.05, ***P<0.001, compared to the control.

### Environmentally or genetically induced hypoxic states mimics loss of *breathless*

The data above indicate that ubiquitous or trachea-specific loss of *btl* induces internal tissue hypoxia, which via reduced insulin leads to slow growth of the entire organism. To confirm that hypoxia *per se* is the cause of this growth inhibition, and to assess whether this inhibition is part of an adaptive response to hypoxia or a pathological result of low physiological oxygen levels, we investigated the effects of environmentally imposed hypoxia on larval growth. Wild-type (*w^1118^*) animals reared under hypoxia (5% O_2_) displayed reduced pupal size, despite a developmental delay of pupariation extending the larval growth period, indicating strong growth inhibition phenocopying that seen with *btl>btl-RNAi* (Fig. 5A, 5B, and S4A).

The signaling pathway that detects low oxygen levels and triggers cellular adaptation programs is conserved between *Drosophila* and mammals. The main effector protein of this pathway is the transcription factor HIF-1a, encoded in *Drosophila* by the gene *similar* (*sima*)^52, 53^. Under normoxic conditions, this protein is marked for proteasomal degradation through the oxygen-dependent activity of HIF-1a prolyl hydroxylase (Hph), also called Fatiga (Fga) in *Drosophila*^54–56^. To differentiate between hypoxia-induced damage versus a genetic adaptive response, we activated the hypoxia-adaptation program by mutation of *Hph*, thus de-repressing HIF-1a-dependent downstream effects under normoxic conditions. Similar to hypoxic conditions or loss of *btl* function, lack of *Hph* resulted in growth inhibition and developmental delay despite the animals’ normoxic environment (Fig. 5A, 5B, and S4A). This confirms that the oxygen-dependent growth effect is under genetic control, rather than arising from a lack of energy due to insufficient aerobic respiration or other pathological damage.

To evaluate whether the specific insulin-related phenotypes seen in *btl*-knockdown animals are components of the genetic hypoxia-adaptation program, we measured *Dilp* expression and DILP retention within the IPCs in hypoxic wild-type (*w^1118^*) animals and in normoxic *Hph* mutants experiencing “genetic hypoxia”. Like knockdown of *btl*, both manipulations led to strongly reduced *Dilp3* and *Dilp5* expression compared to normoxic controls (Fig. 5C). Furthermore, IPC DILP2 and DILP5 levels were increased in hypoxic and *Hph*-mutant animals, indicating that insulin is retained in IPCs (Fig. 5D). The lower effect on DILP3 levels possibly reflects the reduced expression of *Dilp3* (a decrease of up to ~90%; Fig. 5C), which indicates that the observed DILP3 level may in fact represent retention of this DILP as well.

Consistent with decreased systemic insulin signaling, both the low-O_2_ environment and genetic “hypoxia” caused by lack of *Hph* led to reduced pAkt levels (Fig. 5F). To further investigate this, we made use of an *in-vivo* insulin-pathway reporter that reflects intracellular insulin-pathway activity based on plasma-membrane localization of GFP^47^. Under normoxic conditions, GFP fluorescence was localized to the membrane of fat-body cells, indicating high levels of insulin-pathway activity, while under external hypoxia, the reporter remained cytoplasmic, confirming a reduction in insulin signaling (Fig. 5G). Thus, tissue hypoxia, arising from tracheation defects or low external O_2_ levels, leads to the activation of growth-limiting mechanisms downstream of the HIF-1a oxygen-sensing pathway. Activation of this pathway naturally or by genetic manipulation leads to reduced insulin expression and release, reducing systemic insulin signaling and growth. Like animals with *btl* knockdown, hypoxic wild-types and normoxic *Hph* mutants exhibited reduced ecdysone signaling (measured as expression of *E75B*) and upregulation of *bnl* (Fig. S4B and S4C), consistent with an underlying defect common to all three manipulations – namely, the activation of the genetic hypoxia-adaptation program.

**Fig. 5.**
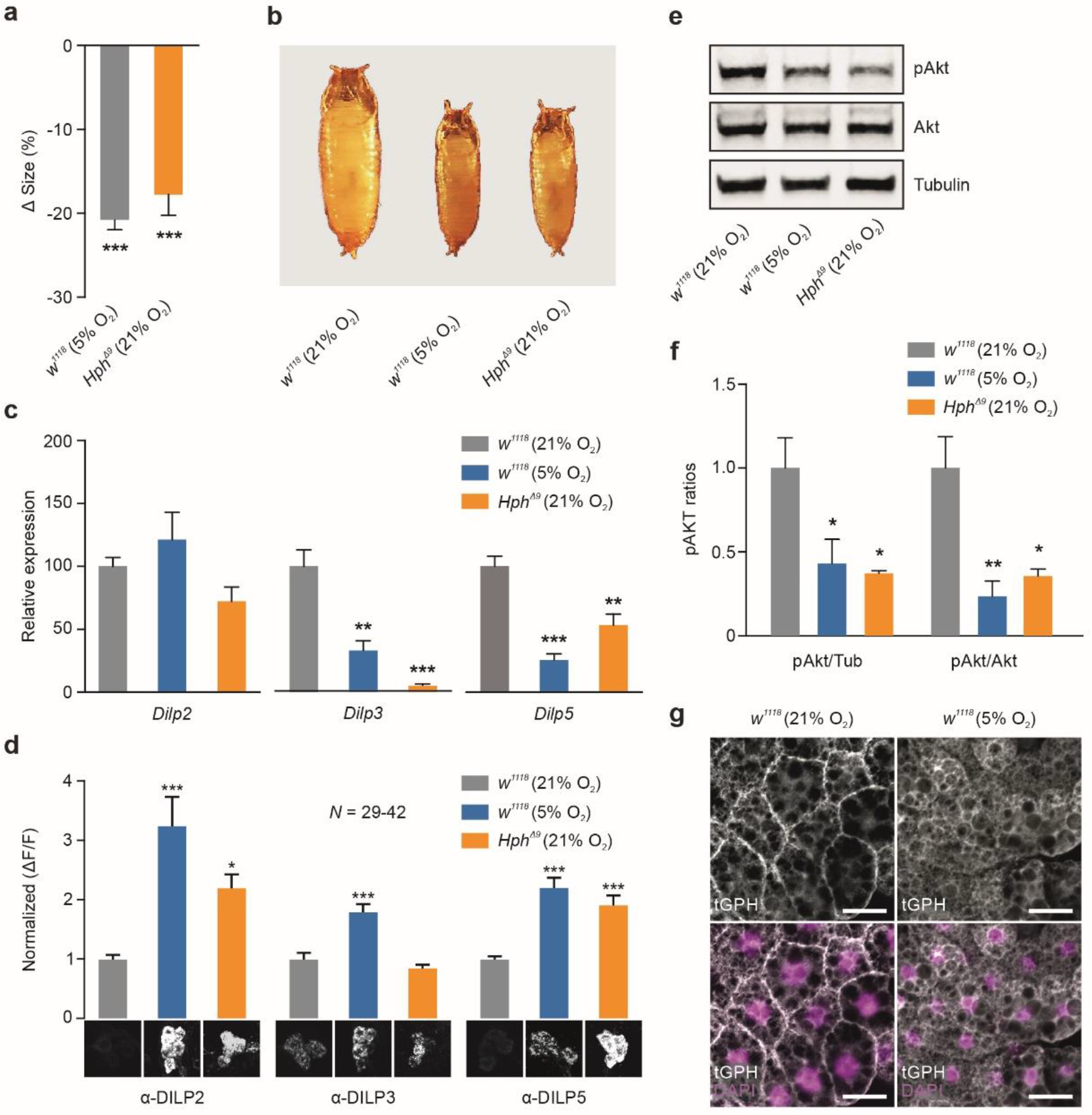
Hypoxia and genetic defects that mimic this condition inhibit growth and insulin secretion. **a-b** Quantification of pupal size (**a**) and representative images (**b**) of animals exposed to hypoxia (5% O_2_) or mutation of *HIF-1a prolyl-hydroxylase* (*Hph*), which genetically mimics hypoxia (even under normoxia conditions, 21% O_2_) compared to controls (*w^1118^* in normoxia). Values are percent change in pupal size versus *w^1118^* in normoxia controls. **c-d** Transcript levels of *Dilp* genes in whole animals (**c**) and DILP peptide levels (**d**) in the IPCs of animals exposed to hypoxia and *Hph* mutants compared to normoxic controls. Representative images of DILP2, −3, and −5 immunostainings in the insulin-producing cells (IPCs) are shown below. **e** Immunoblotting shows that levels of phosphorylated Akt (pAkt) is reduced under hypoxia and in *Hph* mutants compared to *btl>+* controls, quantified in (**f**) from three independent replicates times three larvae each and normalized to Tubulin (Tub) or total Akt levels. Akt and Tubulin were used as loading controls. **g** A transgenic tGPH reporter of insulin signaling confirms that hypoxia reduces systemic insulin signaling in peripheral tissues, represented here by the fat body. Under conditions of normal insulin activity such as normoxia (left), the tGPH sensor is localized to the plasma membrane, while under conditions of low signaling, such as hypoxia, the sensor becomes primarily cytoplasmic. Scale bar, 20 μm. Statistics: one-way ANOVA with Dunnett’s multiple-comparisons test. *P<0.05, **P<0.01, ***P<0.001, compared to the control.

### Loss of *btl* and reduced O_2_ levels cause adipose tissue hypoxia

The fat body is reported to secrete factors into the hemolymph that modulate insulin expression and release from the IPCs^11–13, 15, 16, 57^. To investigate whether this tissue also is involved in the insulinostatic hypoxia-adaptation response, we measured fat-body expression of *bnl* to determine whether this tissue experiences hypoxia under *btl* knockdown in the tracheae. Fat-body *bnl* expression was indeed elevated in these animals, indicating that this tissue experiences hypoxia when Btl-mediated tracheal growth is reduced (Fig. 6A). To confirm the hypoxic state of the fat tissue, we made use of a genetic hypoxia reporter based on a GFP fusion with the oxygen-dependent degradation (ODD) domain of HIF-1a^58^. Under normoxic conditions, GFP::ODD is ubiquitinylated through the activity of Hph, targeting it for degradation. RFP expressed from the same regulatory sequences, but lacking the ODD, acts as a ratiometric control. In the fat body, the ODD::GFP reporter was rapidly degraded under normoxia, leading to a low GFP:RFP ratio (Fig. 6B and 6C). In contrast, an increased ratio of GFP to RFP was observed in animals reared under 5% O_2_, indicating reduced oxygen-dependent degradation of the ODD-linked GFP. Together these data show that the fat body experiences biologically significant hypoxia under genetically or environmentally induced reduced-oxygen conditions. Interestingly, we also observed a dramatic change in the size and quantity of lipid storage droplets in the fat body under hypoxic conditions induced by loss of *btl* (Fig. S5). The fat body is known to release large amounts of lipids under certain conditions. Although the known humoral factors from the fat that affect IPC physiology are peptides and proteins, other classes of signal molecules such as lipids may also be released from the fat under hypoxia to affect insulin secretion.

### A fat-body-derived inhibitory signal represses insulin secretion under hypoxia

We next asked whether the brain and the IPCs alone could sense hypoxia and induce changes in insulin physiology, or whether the fat body might be necessary for organismal growth adaptation to hypoxia. To investigate this, we cultured larval brains (containing IPCs) alone or with fat bodies for 16 hours *ex vivo* under normoxic and hypoxic conditions and determined the effects of these manipulations on insulin expression and secretion (Fig. 6D). While intact hypoxic larvae reduce *Dilp3* and −*5* expression (Fig. 5C), we observed no decrease in expression of the three insulin genes in brains cultured alone under hypoxia compared to normoxia (Fig. 6E). On the other hand, brains co-cultured with fat bodies under hypoxia displayed reduced *Dilp* expression compared to normoxic co-cultures. Together these observations demonstrate that the brain alone is insufficient for the hypoxia-adaptation response, and that a humoral signal from the fat body is required to downregulate *Dilp3* and −*5* expression To confirm these results in whole animals, we blocked the genetic hypoxia-response program in the fat body by expressing fat-body-specific RNAi against *sima/HIF-1a*. In these animals, even when grown under hypoxic conditions, the fat body should behave as if it is normoxic, allowing the contribution of the fat body to organismal responses to be isolated. Animals with fat-body knockdown of *sima/HIF-1a* also fail to downregulate *Dilp* transcription under hypoxia (Fig. 6F). This suggests that Sima/HIF-1a activation in the fat tissue under hypoxic conditions is necessary for the repression of *Dilp* expression in the brain. Furthermore, knockdown of *sima/HIF-1a* in the fat body also blocked the ability of hypoxic animals to repress insulin secretion (Fig. 6G), indicating that activation of HIF-1a is required in the fat body for the hypoxia-adaptation program that modulates IPC insulin secretion.

To confirm the requirement for the presence of the fat body for DILP2 retention under hypoxia, we cultured wild-type brains either alone or with wild-type or *Hph*-mutant fat bodies, under normoxia and hypoxia. DILP2 was retained when brains were co-cultured with wild-type fat-body tissue under hypoxia, but not in brains cultured alone under these conditions (Fig. 6H), indicating again that one or more signals secreted from the fat tissue is required to induce the insulin-retention under hypoxia. To assess the genetic control of this signal, we co-cultured wild-type brains with *Hph*-mutant fat bodies under normoxia. We found that DILP2 was retained in the IPCs, similar to the effect seen in hypoxic co-culture experiments. Taken together these results demonstrate that the activity of the HIF-1a hypoxia-adaptation program within the fat body is the determinant of IPC insulin retention.

**Fig. 6.**
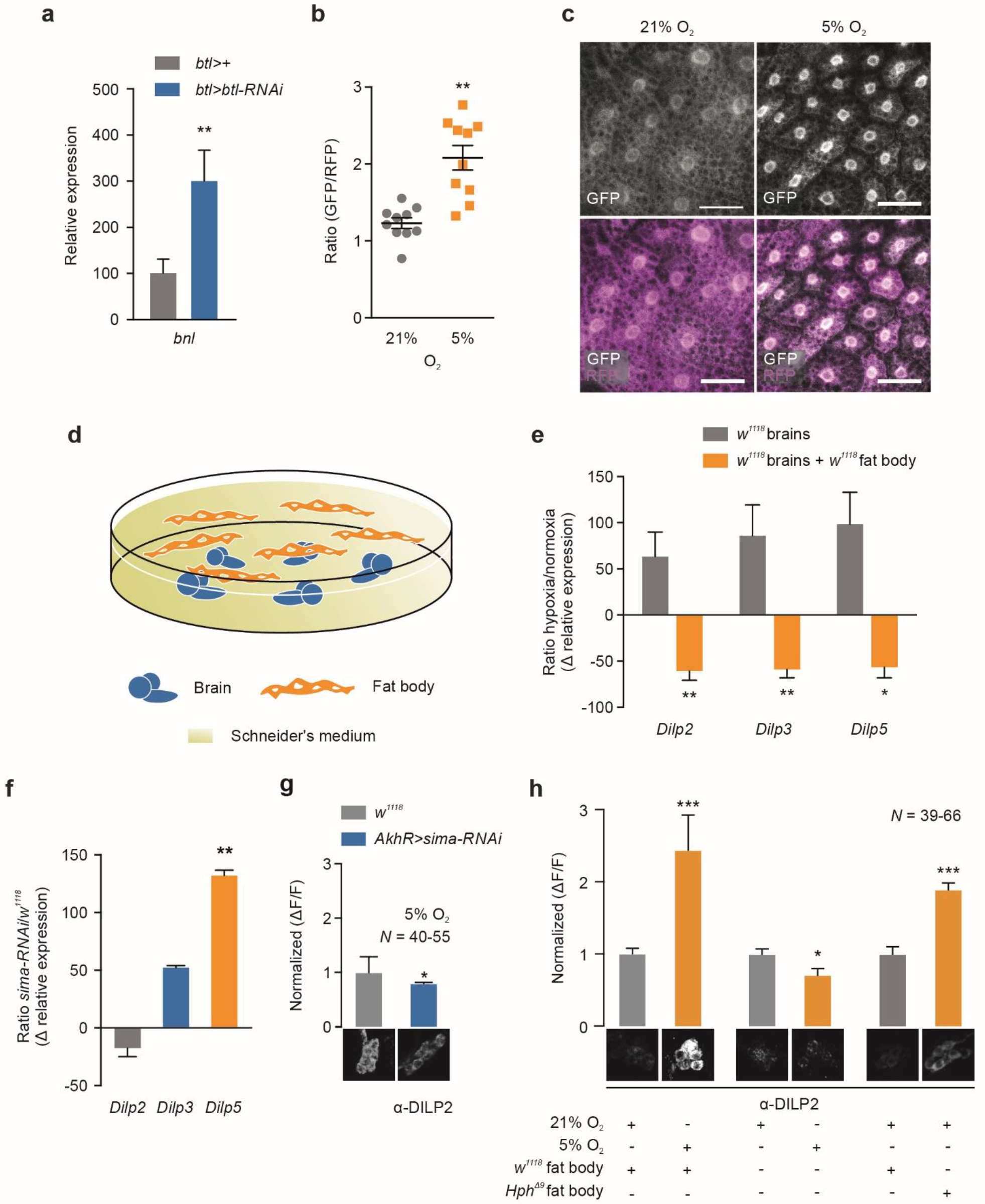
Activation of a fat-body oxygen sensor represses insulin secretion via inter-organ communication. **a** Transcriptional upregulation of the hypoxia-inducible FGF ligand *branchless* (*bnl*) in the fat body of animals with trachea-specific *btl* knockdown indicates hypoxia in this tissue. **b-c** A transgenic reporter of hypoxia indicates that the fat body experiences low oxygen levels that activate HIF-1a when animals are incubated under 5% O_2_ compared to 21% O_2_ conditions. Ratio of integrated GFP (carrying the oxygen-dependent degradation domain of HIF-1a) and RFP (without this domain) signals in fat body (N=10 confocal stacks per condition) (**b**) and representative images (**c**) of GFP::ODD (white) and RFP (magenta). Scale bar, 20 μm. **d** Illustration of *ex-vivo* organ culture technique. **e** Transcript levels of *Dilp* genes in brains of *ex-vivo* organ cultures under hypoxia, compared to normoxia. Expression of *Dilp2*, −*3*, and −*5* increases under hypoxia compared to normoxia (hypoxia transcript levels were normalized to normoxia levels: 0% change). Expression of the three insulin genes decreases when brains were co-incubated with fat-body tissue, closely mimicking the changes seen in whole animals. Thus, some factor(s) from the fat body is required for wild-type IPC responses to hypoxia. **f** Knockdown of *sima/HIF-1a* in the fat body (*AkhR>sima-RNAi*), mimicking normoxia in this tissue, prevents downregulation of *Dilp5* under hypoxia (transcript levels of *Dilp* genes from *AkhR>sima-RNAi* larvae under hypoxia with fat-body-specific *sima* knockdown were normalized to transcript levels of wild-type *w^1118^* in hypoxia: 0% change). **g** Fat-body-specific *sima/HIF-1a* knockdown (subjective fat-body normoxia) reduces hypoxia-induced DILP2 retention. *AkhR>sima-RNAi* larvae exhibit reduced DILP2 accumulation under hypoxia in compared to wild-type *w^1118^* under hypoxia. Representative images of DILP2 immunostainings in the IPCs are shown below. **h** *Ex vivo* co-culture of wild-type *w^1118^* brains with wild-type fat body causes DILP2 retention under hypoxia (left). Brains incubated alone without fat body are unable to induce DILP2 retention under hypoxia (middle). Co-culture of wild-type brains with *Hph*-mutant fat bodies (with activated hypoxia-adaptation program) induces DILP2 retention under normoxia (right) and mimics that seen in true hypoxia (5% O_2_ levels), indicating that the fat-body-derived factor(s) is regulated by the Sima/HIF-1a hypoxia pathway. Representative images of DILP2 immunostainings in the IPCs are shown below. Statistics: Student’s t-test for pairwise comparisons. *P>0.05, **P>0.01, ***P>0.001, compared to the control.

Of the known fat-body derived factors, Eiger is the only one that is reported to be insulinostatic, although its specific effects do not exactly match those we observed under hypoxia ^16^. In any case, we sought to identify any contribution of Eiger to hypoxia-induced growth reduction by expressing RNAi against it in the fat body or against its receptor, Grindelwald, in the IPCs. We rationalized that if Eiger is involved, knockdown of *Eiger* and *Grindelwald* should provide rescue of the growth reduction observed in animals under hypoxic conditions. However, we found that neither *Eiger* nor *Grindelwald* rescued the body-size reduction, suggesting that they do not play a role in hypoxia-induced growth restriction (Fig. S6). We also assessed other fat-body-derived factors, CCHamide-2, Stunted, and Growth-Blocking Peptides 1 and 2, although these are reported to be growth-promoting factors, to rule out their involvement. None of these factors appear to affect hypoxia-induced growth retardation. Thus, the insulinostatic factor released by the fat body appears to be a previously uncharacterized signal that acts inhibitorily on insulin secretion.

## Discussion

### Central fat-tissue oxygen sensing remotely controls insulin secretion via inter-organ communication

In multicellular organisms, oxygen is essential for development and growth. For organisms to adapt their growth and development to oxygen availability, they must have the ability to sense changes in oxygen levels in their environment and to respond by adjusting their growth accordingly. Exposure to hypoxia limits growth during post-embryonic development in most animal species^27, 28, 59^. This appears to be an adaptive response rather than a simple passive effect of insufficient aerobic respiration, since the oxygen levels that reduce growth in many cases are above those that limit cellular metabolism^27, 28^. In this report, we identify one tissue in particular, the fat tissue, that detects internal oxygen levels and is important for inducing the growth-limiting effects of hypoxia. Our data show that, as an adaptive response to oxygen limitation, the fat tissue releases into the circulation one or more humoral factors that inhibit the expression and secretion of insulin from the brain to reduce systemic growth (Fig. 7).

**Fig. 7.**
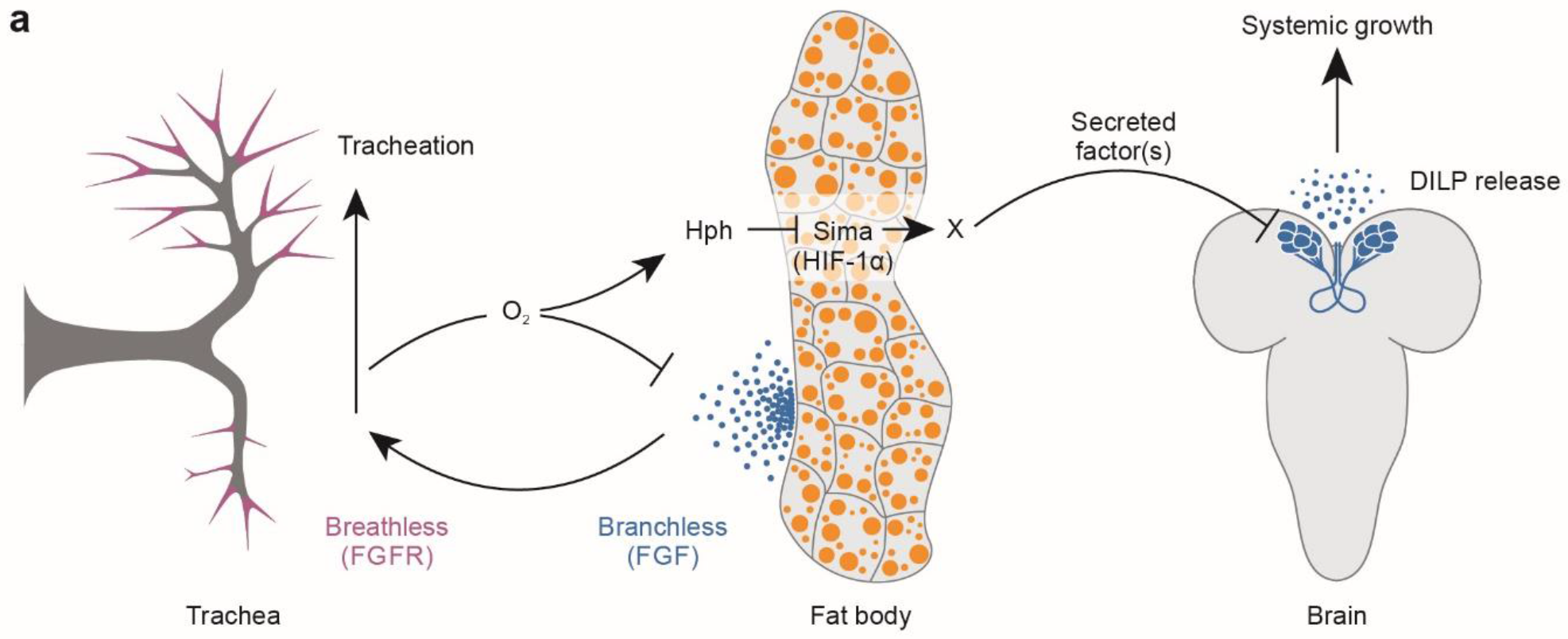
Proposed model for hypoxia sensing and insulin regulation. Tracheation of the fat body is regulated by the FGF ligand Branchless secreted from this tissue in response to local tissue hypoxia, triggering tracheal outgrowth via the FGF receptor Breathless. When the tracheal system is unable to deliver sufficient oxygen because of environmental oxygen levels or body growth, Hph is unable to trigger the degradation of Sima/HIF-1a, which induces the expression or release of one or more humoral factors from the fat tissue that act on the insulin-producing cells (IPCs) to down-regulate *Dilp* transcription and to block DILP release. Hypoxia thereby reduces circulating insulin through activation of a central oxygen sensor in the fat tissue, which slows the growth of the whole organism. During these low-oxygen conditions, activation of Branchless promotes insulin-independent growth of the tracheal airway tubes to increase oxygen supply to internal tissues. This system allows the organism to adapt its metabolism and growth to limited-oxygen conditions by reducing overall body growth through downregulation of insulin signaling while at the same time promoting development of the tracheal system to maintain oxygen homeostasis.

Oxygen homeostasis also requires the coordination of growth between the tissues that deliver oxygen and those that consume it. The development of the oxygen-delivery system is therefore oxygen-sensitive in both mammals and *Drosophila*. In mammals, local tissue hypoxia promotes angiogenesis via induction of many pro-angiogenic factors, including FGF^60^. In *Drosophila*, tissue hypoxia induces expression of the FGF-like ligand *bnl*, leading to branching of the tracheal airway tubes towards oxygen-depleted areas to promote survival by increasing oxygen delivery^33, 61^. Our study shows that this hypoxia-induced fat body mechanism operates downstream of insulin, since reduced insulin signaling in the trachea has no effect of overall body growth. This system therefore allows an adaptive response to low oxygen by reducing overall body-growth via suppression insulin signaling, while promoting hypoxia-induced FGF-dependent tracheal growth to increase oxygen supply to internal tissues.

### Does oxygen transport determine insect size?

Most developing organisms stop growing after reaching a genetically predetermined size that is characteristic for the species. Much of the recent insight into the mechanisms that regulate body size has come from genetic studies in *Drosophila* showing that nutrition regulates body size via signals from the fat body^2, 7, 14–16, 62^. Although this helps explain how organisms modulate their growth and size according to nutritional conditions, the size-sensing mechanism that allows organisms to assess their size and stop their growth when they have reached optimal size has remained elusive. However, recent evidence suggests that body size in insects may be determined by a mechanism that involves oxygen sensing^4^, and oxygen availability is known to place limits on insect body size^20, 22–24^. According to this recent insight, the limited growth ability of the tracheal system during development may limit overall body size due to downstream oxygen sensing^4^. The size of the tracheal system is established at the beginning of each developmental stage and remains largely fixed, aside from terminal branching, as the body grows until it eventually reaches the limit of the system’s ability to deliver oxygen. This allows the body to “assess” its size by sensing internal oxygen concentrations and to terminate growth at a characteristic size that is determined by the size of the tracheal system. Our RNAi screen of the secretome and receptome suggests that the FGF receptor Btl, which is a key factor essential for tracheal growth during development, is a main determinant of body size. Limiting the growth of the tracheal system via loss of *btl* reduces overall growth and body size. Our data therefore support the notion that the tracheal system is part of a size-sensing mechanism that limits growth of the entire body and determines final body size.

### Oxygen-dependent regulation of insulin gene expression via a fat-tissue signal(s)

Our results show that hypoxic conditions and loss of Hph activity selectively repress *Dilp3* and *Dilp5* transcription, while having little or no suppressive effect on *Dilp2* expression. This suggests that transcriptional regulation specifically of *Dilp3* and *Dilp5* is an important component of the response to hypoxia. Consistent with this observation, previous studies have shown that the transcription of *Dilp2*, −*3*, and −*5* are independently regulated^10^. Nutrient deprivation reduces expression of *Dilp3* and −*5* while having no effect on *Dilp2* expression, similar to the effects of exposure to hypoxia. This may favor a model in which the oxygen-sensing mechanism in the fat body converges on the nutrient sensor via inhibition of TOR. However, it is also possible that *Dilp3* and −*5* are regulated via a fat-body HIF-1a-dependent mechanism that is independent of TOR and that these two *Dilp*s generally show a greater transcriptional response compared to *Dilp2*. This would also be consistent with the notion that one or more novel insulinostatic adipokine factors are involved in mediating the response, since Eiger, which inhibits insulin downstream of the TOR nutrient sensor, regulates *Dilp2* and −*5*, but not *Dilp3*^16^, which we show here is the main *Dilp* gene responding transcriptionally to low-oxygen conditions.

The transcriptional response could also conceivably be a result of autocrine feedback regulation that regulates insulin in both *Drosophila* and mammals^63, 64^. However, lower secretion of DILPs, as we observe under hypoxic conditions, is generally associated with increased expression of *Dilp3* and −*5*, which suggests that the hypoxia-induced repression of *Dilp3* and −*5* is a specific transcriptional response and not a consequence of autocrine effects. Together our results indicate that part of oxygen-dependent growth regulation is mediated by transcriptional control of *Dilp3* and −*5*. It will be important to identify in future studies the fat-tissue factor(s) involved and the mechanism by which insulin production is regulated.

### Effects of tissue hypoxia in mammals

Cell and tissue hypoxia are also observed in human conditions of obesity and cancer. The insect fat body performs the functions of mammalian fat and liver tissues, serving as an endocrine organ that secretes factors that affect insulin signaling^11, 12, 14–16, 65^. Accordingly, perturbation of insulin signaling by adipose and hepatic tissue hypoxia is also observed in mammalian systems^66^. In mammals, obesity induces hypoxia within adipose tissue due to the rarefaction of vascularization of this tissue and the resulting low blood flow^67^. This leads to the production and release of inflammatory mediators and other adipokines that are associated with the pathophysiology of obesity-related metabolic disorders including diabetes^68–73^. Although loss of normal β-cell activity is considered a main factor in diabetes, the mechanism by which tissue hypoxia affects insulin secretion is poorly understood. Our finding of one or more hypoxia-induced fat-body-derived insulinostatic humoral factors opens new avenues for understanding the role of adipose-tissue hypoxia in obesity and its impact on diabetes.

Obesity also leads to physical and hormonal changes that affect breathing patterns, causing apnea and thus intermittent episodes of systemic hypoxia^74^. These hypoxic periods can induce changes in the liver, leading to fatty liver disease and dyslipidemia, which have their own insulinotropic effects. Furthermore, hypoxia-induced programs play important roles in tumor formation. During cancer development, tumor cells undergo a metabolic reprogramming, the so-called “Warburg effect,” in which their metabolism shifts from oxidative phosphorylation to glycolysis^75^. Activation of HIF-1a is believed to play a key role in driving this shift. Because the hypoxia-sensing mechanism and the insulin-signaling system are conserved between flies and mammals, understanding the effects of hypoxia on the fat body could thus provide insight into many human disease states.

### The identity of the oxygen-dependent adipokine(s) that inhibits insulin secretion

Our work shows that one or more insulinostatic factors relay information from the fat-body oxygen sensor to the IPCs about the amount of internal oxygen to adapt growth and metabolism to oxygen conditions. This raises the question of the identity and function of the humoral signal(s). Many of the known *Drosophila* adipokines that affect insulin secretion from the IPCs are regulated by nutrient-dependent TOR-pathway activity in the fat body, including CCHamide-2^12^, Eiger^16^, FIT^13^, GBP1 and GBP2^14^, and Stunted^15^. On the other hand, our data suggest that the humoral factor(s) released by the fat under hypoxia is released in response to hypoxia-induced HIF-1a pathway activation. However, hypoxia is known to inhibit TOR pathway activity via Sima/HIF-1a activation of the TSC1/2 activator Scylla/REDD1^76, 77^, making it possible that the HIF-1a-dependent fat-body oxygen-sensing mechanism converges on the TOR-dependent nutrient-sensing mechanism. Since our *ex-vivo* organ-culture experiments demonstrate that the signal(s) released by the fat body under hypoxia is inhibitory, Eiger (orthologous to the pro-inflammatory cytokine TNF-alpha), which inhibits *Dilp-*gene expression, seems the most apt candidate among the known factors. However, hypoxia strongly induces DILP retention, while Eiger is reported to have no effect on this process, and Eiger does not affect *Dilp3* expression^16^. Furthermore, fat-body RNAi against *Eiger* and knockdown of its receptor *Grindelwald* in the IPCs had no effect on hypoxia-induced growth inhibition. Thus, if Eiger is involved at all in mediating the effect of fat-body hypoxia on IPC activity, it likely acts in coordination with other adipokine factors.

In mammals, several adipokines regulate β-cell function and insulin secretion, including leptin, which conveys information about fat storage and is a functional analog of the *Drosophila* fat-derived cytokine Unpaired-2^11, 78^. Interestingly, we observed that hypoxia induced by *btl* knockdown caused strong increase in fat-body lipid-droplet size, indicating a change in lipid metabolism within the fat tissue. It is conceivable that the signal(s) released by the fat body in response to hypoxia is a lipid or a lipid-binding protein. For example, the mammalian Fatty Acid Binding Protein 4 (FABP4) is an insulin-modulating adipokine that is influenced by obesogenic conditions that lead to adipose tissue hypoxia^79^, and orthologous proteins are encoded by the *Drosophila* genome. Although this shows that fatty-acid-binding proteins can regulate insulin secretion, FABP4 stimulates insulin secretion, while our data suggest that the adipokine relaying information between the fat-tissue oxygen sensor and the IPCs acts as an inhibitor of insulin secretion. Our study opens future avenues for uncovering inter-organ crosstalk mechanism that regulate insulin secretion in relation to conditions associated with hypoxia.

In conclusion, our study unravels a mechanism that allows organisms to adapt their metabolism and growth to environments with low oxygen. Hypoxia activates a fat-tissue oxygen sensor that remotely controls the secretion of insulin from the brain by inter-organ communication. This involves the activation of a HIF-1a-dependent genetic hypoxia-adaptation program within the fat tissue, which then secretes one or more humoral signals that alter insulin-gene expression and repress insulin secretion, thereby slowing growth. Given the conservation of oxygen-sensing systems and the influence of oxygen on growth between *Drosophila* and mammals, a similar adaption response may operate in mammals via adipose tissue oxygen-sensing to maintain oxygen homeostasis.

## Methods

### Fly strains and husbandry

Animals were reared on standard cornmeal-yeast medium (Nutri-Fly, Bloomington recipe) at 25° and 60% relative humidity, under a 12L:12D light cycle. The *AkhR-GAL4∷p65* transgene containing the enhanced activator *GAL4∷p65*^80^ drives very strongly in the larval fat body. The *GAL4∷p65* was “recombineered” into genomic BAC construct CH322-147J17^81^ obtained from Children’s Hospital Oakland Research Institute (Oakland, CA, US), replacing the first coding exon, and the construct was integrated into the *attP2* genomic site by standard injection methods. The following stocks were obtained from the Bloomington *Drosophila* stock center: *arm-GAL4* (#1560), *btl-GAL4* (#41803), *da-GAL4* (#55850), *elav-GAL4*^82^ (#458), *Mef2-GAL*4^83^ (#27390), *ppl-GAL4* (#58768), *R96A08-GAL4^84^* (#48030), *tGPH* insulin-activity sensor^47^ (#8164), *UAS-CCHa2-RNAi* (#57183), *UAS-CCHa2-R-RNAi #2* (#25855), *UAS-eiger-RNAi* (#58993), *UAS-InR-RNAi* (#992), *UAS-Luciferase-RNAi* (#35789), *UAS-methuselah-RNAi #1* (#27495) and *#2* (#36823), *UAS-methuselah-like-10-RNAi #1* (#62315) and *#2* (#51753), *UAS-Tor^TED^* (TOR dominant negative; #7013), and *UAS-TSC1/2*^85^. Vienna *Drosophila* Resource Center (VDRC) RNAi lines used for follow-up studies include *UAS-Akt-RNAi* (#103703), *UAS-bnl-RNAi* (#101377), *UAS-btl-RNA^GD950^* (#950), *UAS-btl-RNAi^GD27108^* (#27108), *UAS-CCHa2-R-RNAi #1* (#100290)^12^, *UAS-grindelwald-RNAi #1* (#104538)^16^ and *#2* (#43454)^16^, *UAS-sima-RNAi* (#106187), and *UAS-Stunted-RNAi* (#23685)^15^. *ΔGbp1,2^14^* was kindly given by T. Koyama. *Hph^Δ9^* (*Fatiga^Δ9^*, *Fga^Δ9^*) was a kind gift of P. Wappner, *ubi-ODD∷GFP* hypoxia sensor^58^ was a kind gift of S. Luschnig, and *UAS-FLAG∷Dilp2*^86^ was a gift of E. Hafen and H. Stocker. The common laboratory stock *w^1118^* from VDRC (#60000) was used as a control in many experiments.

### RNAi screen for pupal size

A list of genes encoding the secretome (secreted proteins) and receptome (membrane-associated proteins) was generated using ENSEMBL and GLAD gene-ontology databases^87, 88^. UAS-RNAi lines against 1,845 genes for the screening of the secretome and receptome were obtained from the Bloomington *Drosophila* stock center^36^ and the Vienna *Drosophila* Resource Center (VDRC)^35^, and transformant IDs are given in Table S1. Males from each RNAi line were crossed to *da-GAL4* virgin females, animals were allowed to lay eggs for 24 hours, and the average size of pupal offspring was determined from images acquired with a Point Grey Grasshopper3 camera using a custom MATLAB script (MATLAB R2016b, The MathWorks, Inc., Natick, Massachusetts, USA)^89^. Each line was given a *Z*-score, calculated as the number of SD between that line’s average and the average of all lines. Lines with a *Z*-score between −2 and 2, corresponding in this data set roughly to a ±15% size abnormality, were not analyzed further, while 89 lines with greater effect were identified.

### Tracheal measurements and hypoxic survival rates

To determine tracheal length and branching, L3 larvae [96 hours after egg lay (AEL) for control; 120 hours for *btl-RNAi* animals to account for slowed growth] were heat-fixed in a drop of glycerol on a microscope slide on a 60 °C heat block until they became relaxed. Tracheal cells and branching of the fat-body terminal branches were visualized in whole larvae by imaging of tracheae labeled with GFP or by direct light microscopy. Length and branching were manually quantified using the FIJI (NIH) software package^34, 90^. To determine survival rates of *btl* knockdown animals, timed egg-lays were performed, and first-instar (L1) larvae were transferred to fly vials containing standard food and incubated in either hypoxia (5% O_2_) or normoxia (21% O_2_, normal atmospheric oxygen concentration), and the number of animals surviving to the pupal stage was determined. The low-oxygen environment was generated by slowly passing 5% O_2_, 95% N2 through a clear, airtight plastic chamber placed inside a 25 °C, 12L:12D incubator. Gas was bubbled through water upstream and downstream of the chamber, to provide humidity and to prevent infiltration of outside air, and vented outside the incubator. The chamber was flushed with high-flow-rate gas for 5 minutes after any opening.

### Analysis of developmental timing and growth rates

To synchronize larvae for growth-rate and developmental-timing assays, flies were allowed to lay eggs for 4 hours on apple-juice agar plates supplemented with yeast paste. Newly hatched L1 larvae were transferred into vials with standard food (30 per vial to prevent crowding) 24 hours later. To determine growth rates, synchronized larvae were weighed at time points 75-96 hours AEL in groups of 5-10 animals. For developmental timing experiments, pupariation time was determined manually.

### *Ex-vivo* organ co-culture

For organ co-culture experiments, brains and fat bodies were dissected from 96-hour-AEL wild-type *w^1118^* larvae or 120-hour-AEL *Hph*-mutant larvae (to adjust for their slow growth) in cold Schneider’s culture medium (Sigma-Aldrich #S0146) containing 5% fetal bovine serum (Sigma). Fifty brains, with or without fat bodies present, were incubated in 50 μl culture medium for 16 hours at 25 °C in small dishes in moist chambers in either hypoxia or normoxia. After incubation, brains were removed for immunostaining and qPCR analyses.

### Expression analysis by qPCR

To quantify gene expression by quantitative real-time PCR (qPCR), mRNA was prepared from six sets of five whole larvae (96 hours AEL for *btl>+* controls and normoxia animals, and 120 hours AEL for *btl-RNAi* larvae, animals raised under hypoxia, and *Hph* mutants to adjust for their slower growth) using the RNeasy Mini Kit (Qiagen #74106) with DNase treatment (Qiagen #79254). RNA yield was determined using a NanoDrop spectrophotometer (ThermoFisher Scientific), and cDNA was generated using the High-Capacity cDNA Reverse Transcription Kit with RNase Inhibitor (ThermoFisher #4368814). qPCR was performed using the QuantiTect SYBR Green PCR Kit (Fisher Scientific #204145) and an Mx3005P qPCR System (Agilent Technologies). Expression levels were normalized against *RpL32*. The primer pairs used are listed below.
*bnl:* TGCCCTATCACAGAGTTGC and ACCTACACGAACGCCATCAC
*E75B:* CAACAGCAACAACACCCAGA and CAGATCGGCACATGGCTTT
*Dilp2:* CTCAACGAGGTGCTGAGTATG and GAGTTATCCTCCTCCTCGAACT
*Dilp3:* CAACGCAATGACCAAGAGAAC and GCATCTGAACCGAACTATCACTC
*Dilp5:* ATGGACATGCTGAGGGTTG and GTGGTGAGATTCGGAGCTATC
*InR:* CTCAGCCATACCAGGGACTTT and CTCTCCATAACACCGCCATC
*RpL32:* AGTATCTGATGCCCAACATCG and CAATCTCCTTGCGCTTCTTG

### Western-blotting analysis

To quantify signaling-pathway activity downstream of InR, the level of Akt phosphorylation (pAkt) was determined by Western blot. For hypoxia experiments, larvae were transferred to 5% O_2_ at 72 hours AEL and analyzed 24 hours later, compared with 96-hour-AEL normoxic controls. *Hph* mutant larvae were assayed at 120 hours AEL. For each sample, three larvae were homogenized in SDS loading buffer (Bio-Rad), boiled for five minutes, centrifuged at 14,000×g for five minutes to pellet debris, and electrophoresed through a precast a 4-20% polyacrylamide gradient gel (Bio-Rad). Proteins were transferred to a PVDF membrane (Millipore), and the membrane was blocked with Odyssey Blocking Buffer (LI-COR) and incubated with antibodies against pAkt (Cell Signaling Technology #4054, diluted 1:1000), pan-Akt (Cell Signaling Technology #4691, diluted 1:1000), and α-Tubulin (Sigma #T9026, diluted 1:5000) in Odyssey Blocking Buffer (LI-COR) containing 0.2% Tween 20. Primary antibodies were detected with labeled infrared dye secondary antibodies IRDye 680RD and 800CW diluted 1:10,000 (LI-COR), and bands were visualized using an Odyssey Fc imaging system (LI-COR).

### Immunostaining

Brains were removed from *ex-vivo* cultures or dissected from larvae 96-hour-AEL (normoxia) or 120-hour-AEL (hypoxia and *Hph* mutants) in cold Schneider’s culture medium and fixed in fresh 4% paraformaldehyde in PBS for 1 hour. Tissues were washed four times in PBST (PBS containing 0.1% Triton X-100), blocked with PBST containing 3% normal goat serum (Sigma-Aldrich #G9023) at room temperature for 1 hour, and incubated with primary antibodies overnight at 4 °C. The following primary antibodies were used: rat anti-DILP2^57^ (kind gift of P. Léopold) at 1:250 dilution; mouse anti-DILP3 at 1:200 dilution (kind gift of J. Veenstra); and rabbit anti-DILP5^91^ at 1:2,000 dilution (kind gift of D. Nässel). Tissues were washed three times in PBST and incubated in secondary antibodies: goat anti-rabbit Alexa Fluor 488 (Thermo Fisher Scientific, #A32731), goat anti-rat Alexa Fluor 555 (Thermo Fisher Scientific, #A21434), and goat anti-mouse Alexa Fluor 647 (Thermo Fisher Scientific, #A28181), all diluted 1:500 in PBST, overnight at 4 °C. After four washes in PBST, tissues were mounted in Vectashield (Vector Labs #H-1000) and imaged using a Zeiss LSM 800 confocal microscope and Zen software. Retained insulin levels were calculated using the FIJI software package.

### Fat-body tGPH insulin-sensor, ODD∷GFP oxygen-sensor, and lipid-droplet analysis

A GFP-pleckstrin homology domain fusion protein (GPH) under control of the ubiquitous *β-Tubulin* promoter (tGPH) was used as an *in vivo* indicator of intracellular insulin signal transduction in the fat body. Insulin signaling activity causes recruitment of the GPH protein to the plasma membrane^47^. Hypoxia was induced 72 hours AEL, and hypoxic animals were assayed for fat-body GFP localization 24 hours later together with age-matched 96 hour AEL normoxic controls. Fat bodies were dissected in cold Schneider’s medium and immediately imaged. A GFP∷ODD sensor line, made up of the oxygen-dependent degradation domain of Sima fused to GFP, expressed throughout the animal from the *ubiquitin-69E* promoter, was used to determine whether the fat body experiences biologically significant hypoxia under 5% oxygen conditions^58^. This reporter protein is degraded at a rate proportional to tissue oxygen levels. Unmodified RFP expressed from the same regulatory elements provides a ratiometric denominator. Larvae were reared under normoxia or hypoxia, and at 96 or 120 hours AEL, respectively, fat bodies were dissected in cold Schneider’s medium and immediately imaged. Levels of GFP and RFP signals were integrated using the FIJI package, and the ratio between them was calculated. Lipid droplets were detected using label-free Coherent Anti-Stokes Raman Scattering (CARS) microscopy. Fat bodies were dissected in cold Schneider’s culture medium and immediately imaged using a Leica TCS SP8 confocal microscope as described^92, 93^. Lipid droplet number and density were quantified using FIJI software.

### RNAi mini-screen of fat-body factors and IPC receptors

*R96A08-GAL4* targeting the IPCs was crossed to RNAi constructs against receptors, and the strong fat-body driver *AkhR-GAL4∷p65* described above was crossed to lines targeting secreted factors. Newly hatched larvae from 4-hour egg lays were transferred at 24 hours AEL to standard food vials and incubated at 25 °C in normoxia or hypoxia (5% O_2_) until pupariation. Pupal size was measured and for each genotype, the ratio for each possible hypoxic and normoxic vial pair was computed to make up each data set. Experimental data were compared to data from controls expressing *UAS-Luciferase-RNAi*, except for the *ΔGbp1,2* deletion line, which was normalized to *w^1118^*.

### Statistical analyses

Statistical tests and P-values are given within each figure legend. Statistics were calculated using GraphPad PRISM Software, applying ANOVA and Dunnett’s test for multiple-comparisons tests and two-tailed Student’s *t*-tests for pair-wise comparisons. Error bars indicate standard error of the mean (SEM), and P-values are indicated as: *, P<0.05; **, P<0.01; and ***, P<0.001.

**Fig. S1.**
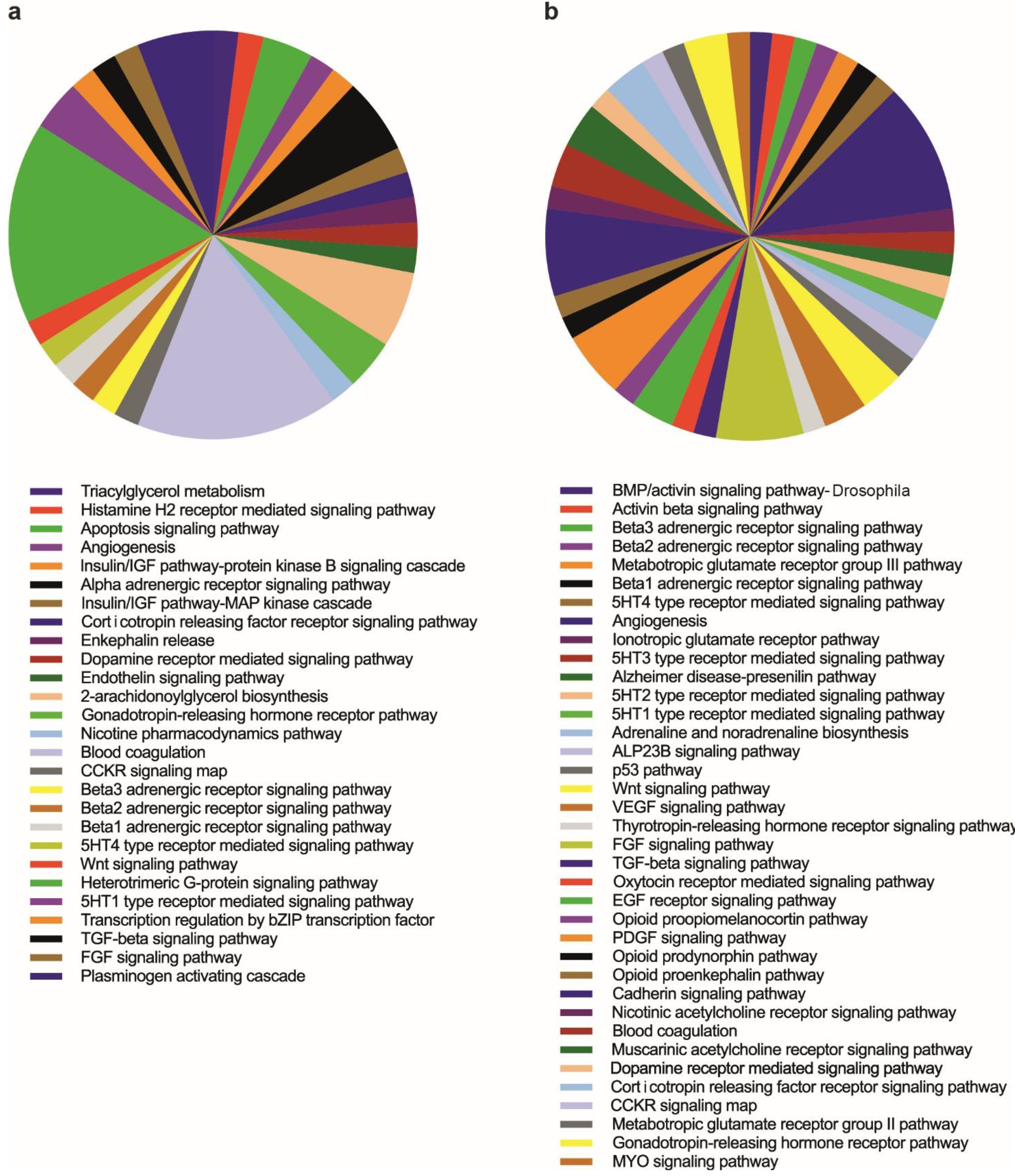
**a-b** Gene-ontology (GO) analysis of hits from the RNAi whose knockdown is associated with increased (**a**) or decreased (**b**) pupal size.

**Fig. S2.**
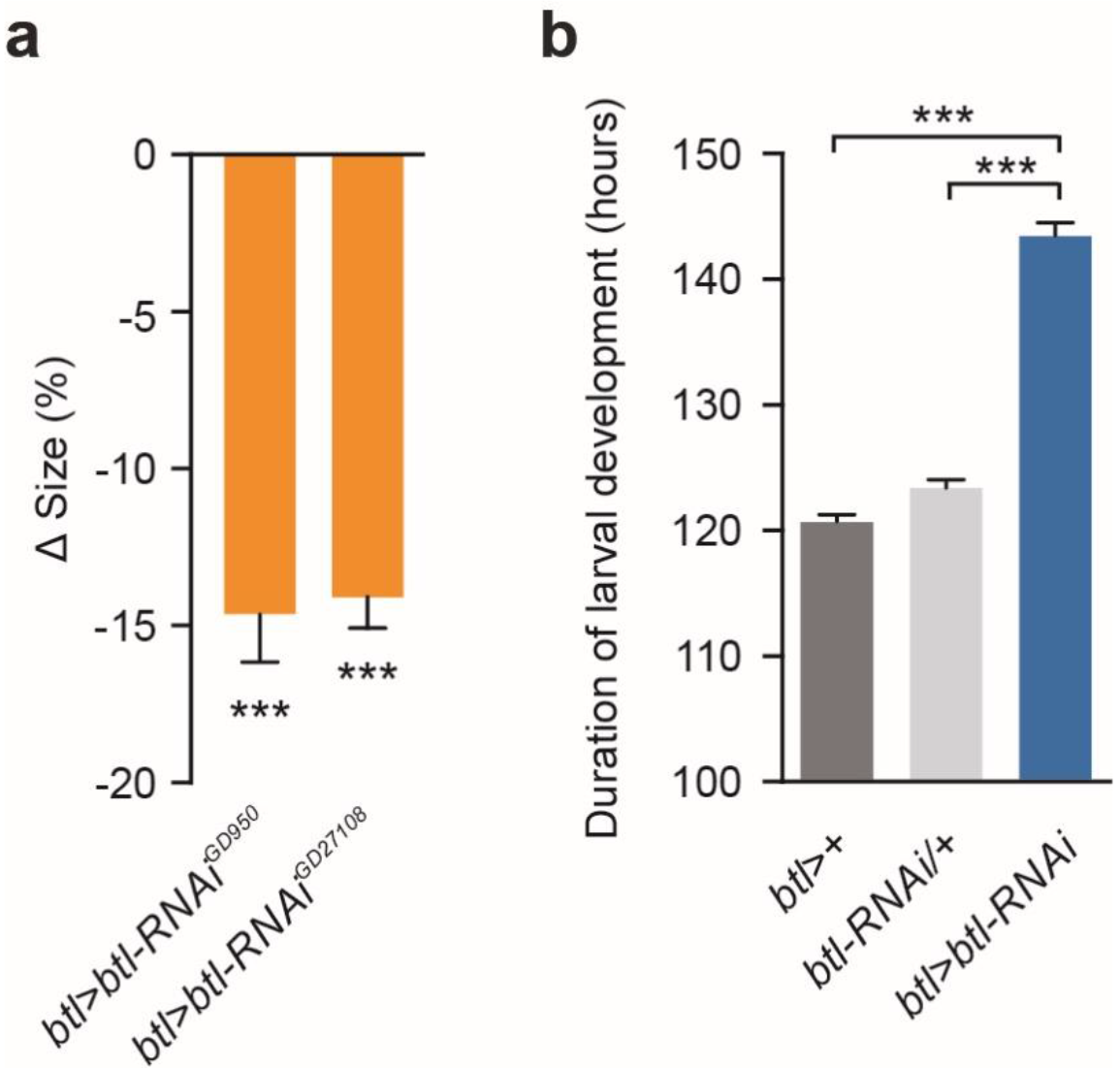
**a** Knockdown of *breathless* (*btl*) in the trachea (*btl>btl-RNAi*) with two additional RNAi lines from VDRC (transformant ID number 950 and 27108) targeting independent sequences of *btl* reduces pupal body size compared to the control (*btl*> crossed to wild type, *w^1118^*). **b** Duration of larval development determined by the onset of pupariation of animals with trachea-specific *btl* knockdown compared *btl>+* driver (*btl>* crossed to wild type, *w^1118^*) and *btl-RNAi/+* (*btl-RNAi* crossed to wild type, *w^1118^*) controls. Statistics: one-way ANOVA with Dunnett’s multiple-comparisons test. ***P<0.001, compared to the control.

**Fig. S3.**
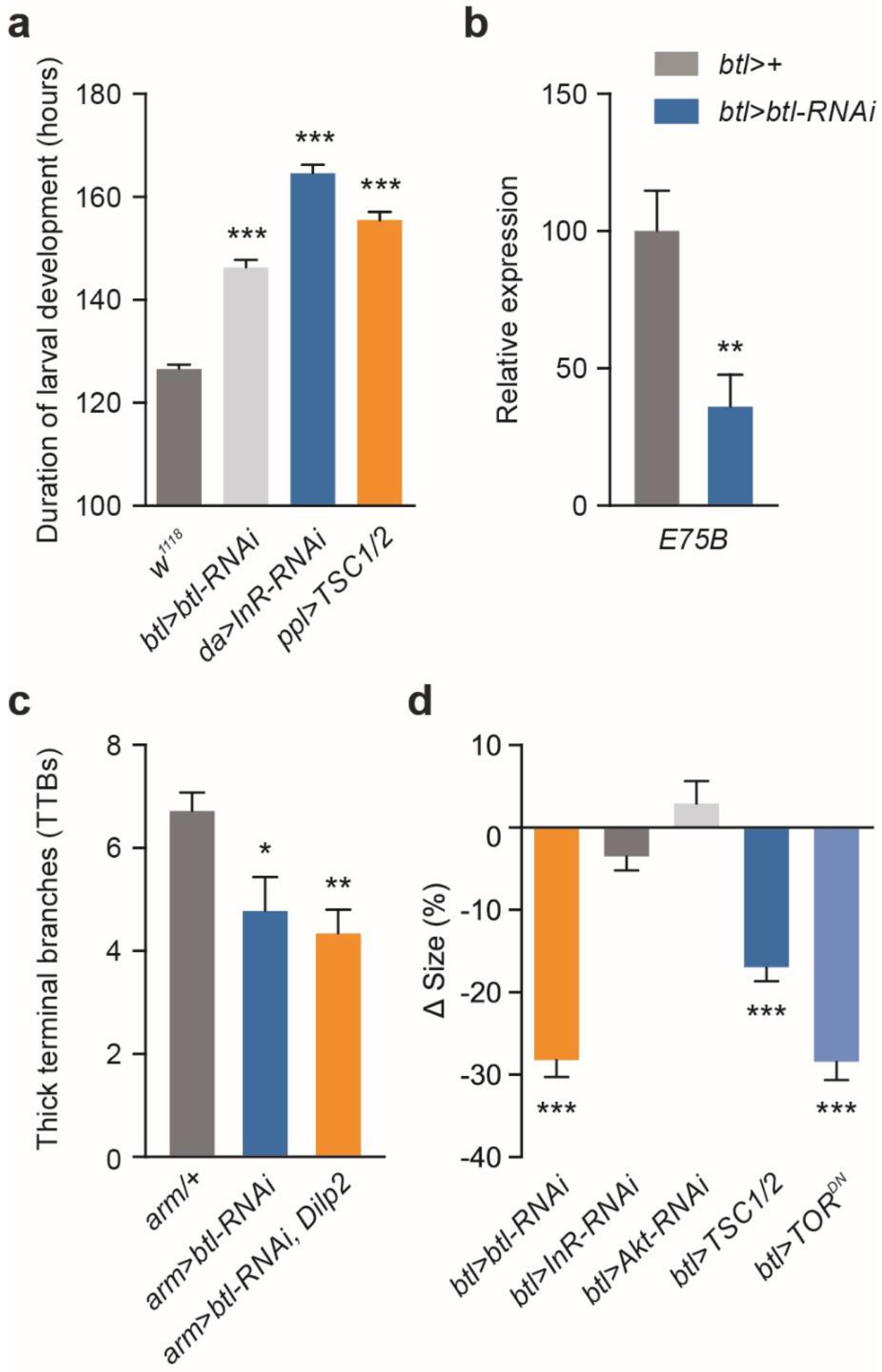
**a** Duration of larval development determined by the onset of pupariation of *da>InR-RNAi* animals with global knockdown of the insulin receptor (*InR*), and *ppl>TSC1/2* animals with fat-body-specific TOR inhibition compared to controls (*w^1118^*). **b** Transcript levels of the ecdysone-inducible *E75B* gene. **c** Effect of global *btl* knockdown and *Dilp2* overexpression using the weak ubiquitous *arm>* driver on the number of thick terminal branches (TTBs) per terminal cell, quantified as the number of cell projections. **d** Pupal size changes in animals with trachea-specific loss of *breathless (btl)* (*btl>btl-RNAi*), insulin signal transduction (*btl>InR-RNAi* and *btl>AKT-RNAi*), or TOR signaling (*btl>TSC1/2* and *btl>TOR^DN^*) compared to *btl>+* controls (*btl>* crossed to wild type, *w^1118^*). Statistics: one-way ANOVA with Dunnett’s test for multiple-comparisons and Student’s t-test for pairwise comparisons. *P<0.05, **P<0.01, ***P<0.001, compared to the control.

**Fig. S4.**
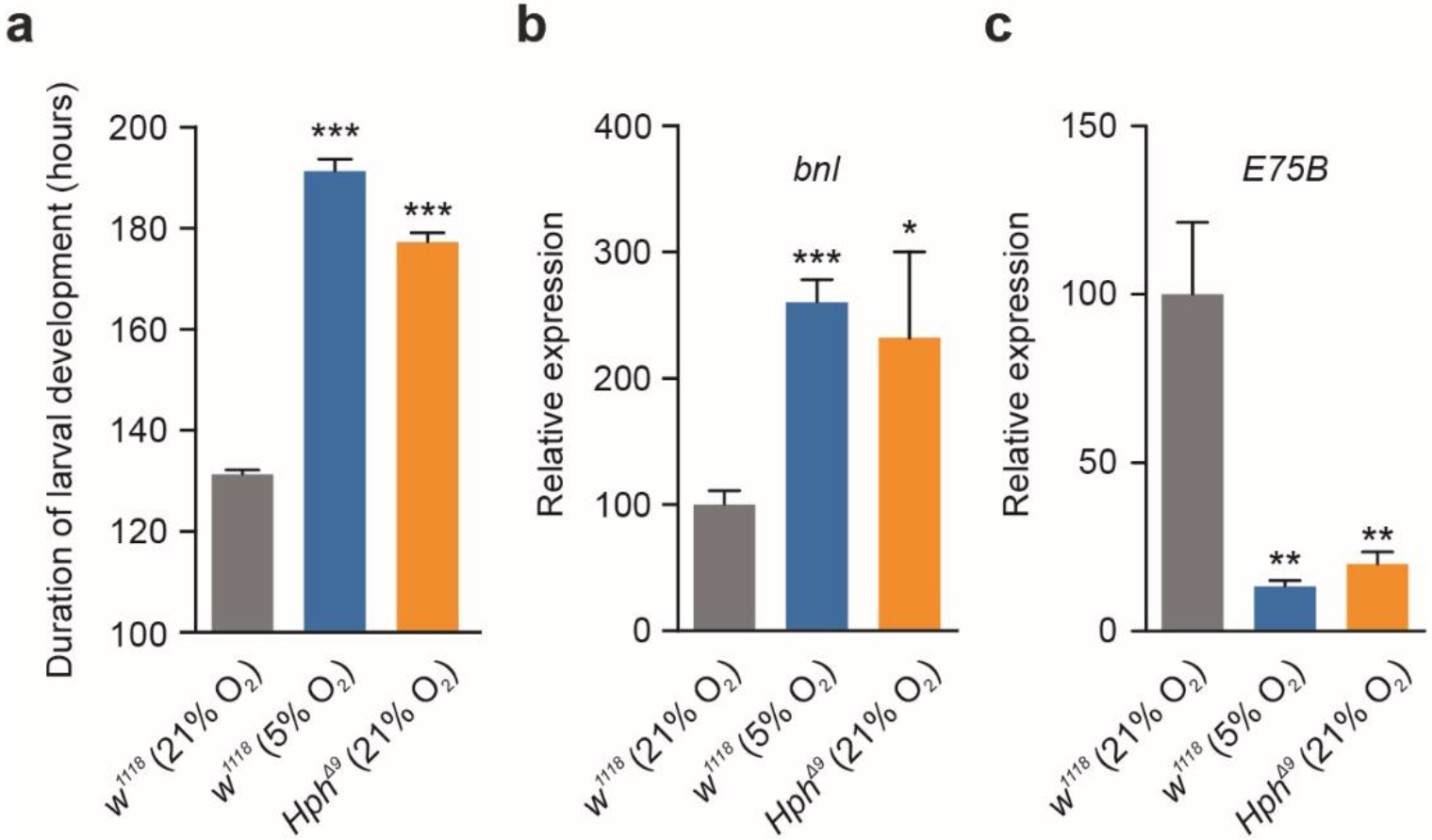
**a** Duration of larval development determined by the onset of pupariation of wild-type (*w^1118^*) animals under normoxia (21% O_2_) and hypoxia (5% O_2_), and *Hph* mutant animals under normoxia. **b-c** Transcript levels of *branchless* (*bnl*) (**b**) and ecdysone-inducible gene *E75B* (**c**) in whole larvae under normoxia in wild-types and *Hph* mutants and in wild-types under hypoxic conditions. Statistics: one-way ANOVA with Dunnett’s multiple-comparisons test. *P<0.05, **P<0.01, ***P<0.001, compared to the control.

**Fig. S5.**
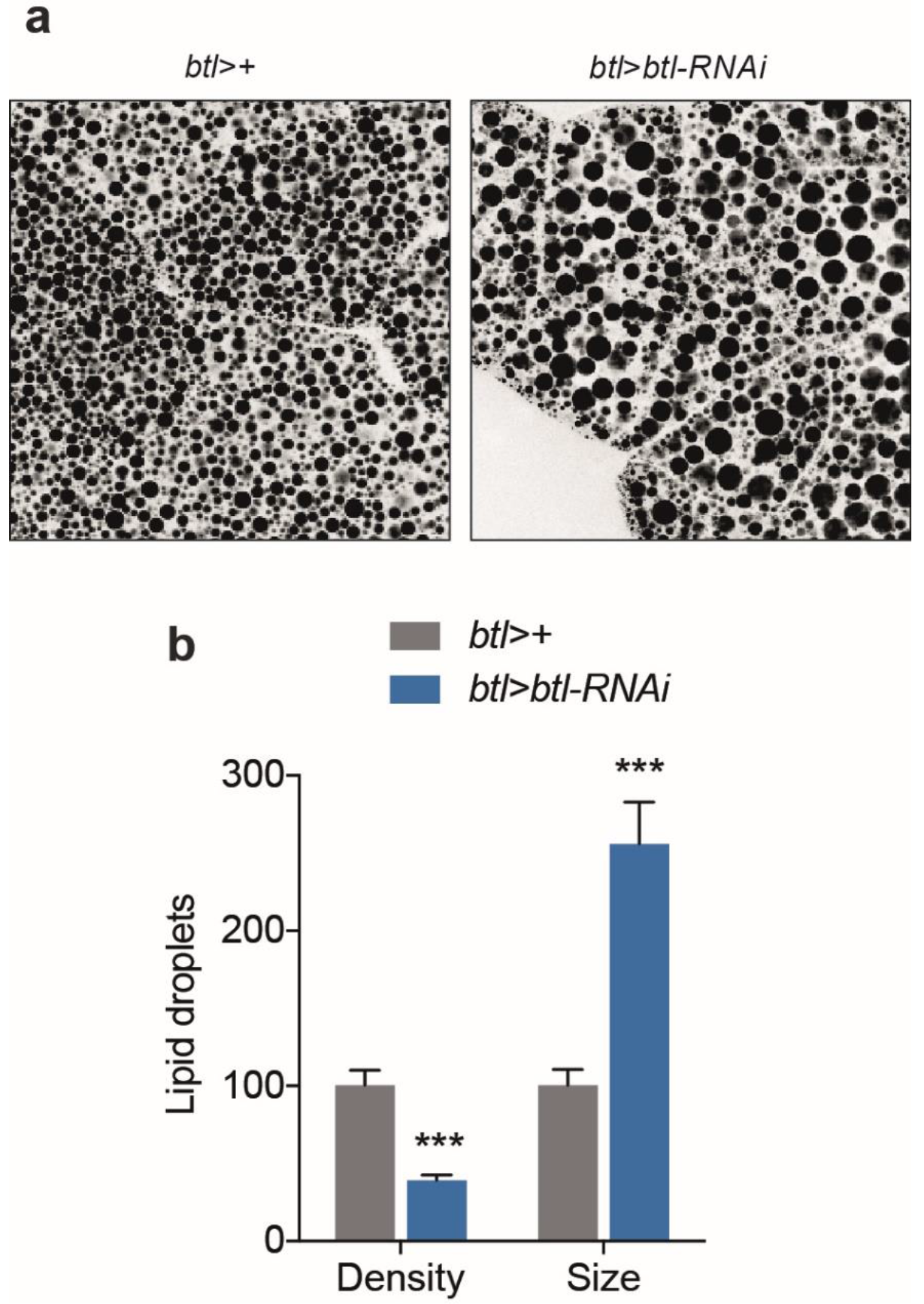
Hypoxia induced by loss of *btl* causes alteration in lipid distribution in the fat body. **a -b** representative images (**a**) and quantification of density and size (**b**) of lipid storage droplets in the fat body detected by CARS microscopy. Controls; *btl>+* (*btl>* crossed to wild type, *w^1118^*). Statistics: Student’s t-test for pairwise comparisons. ***P<0.001, compared to the control.

**Fig. S6.**
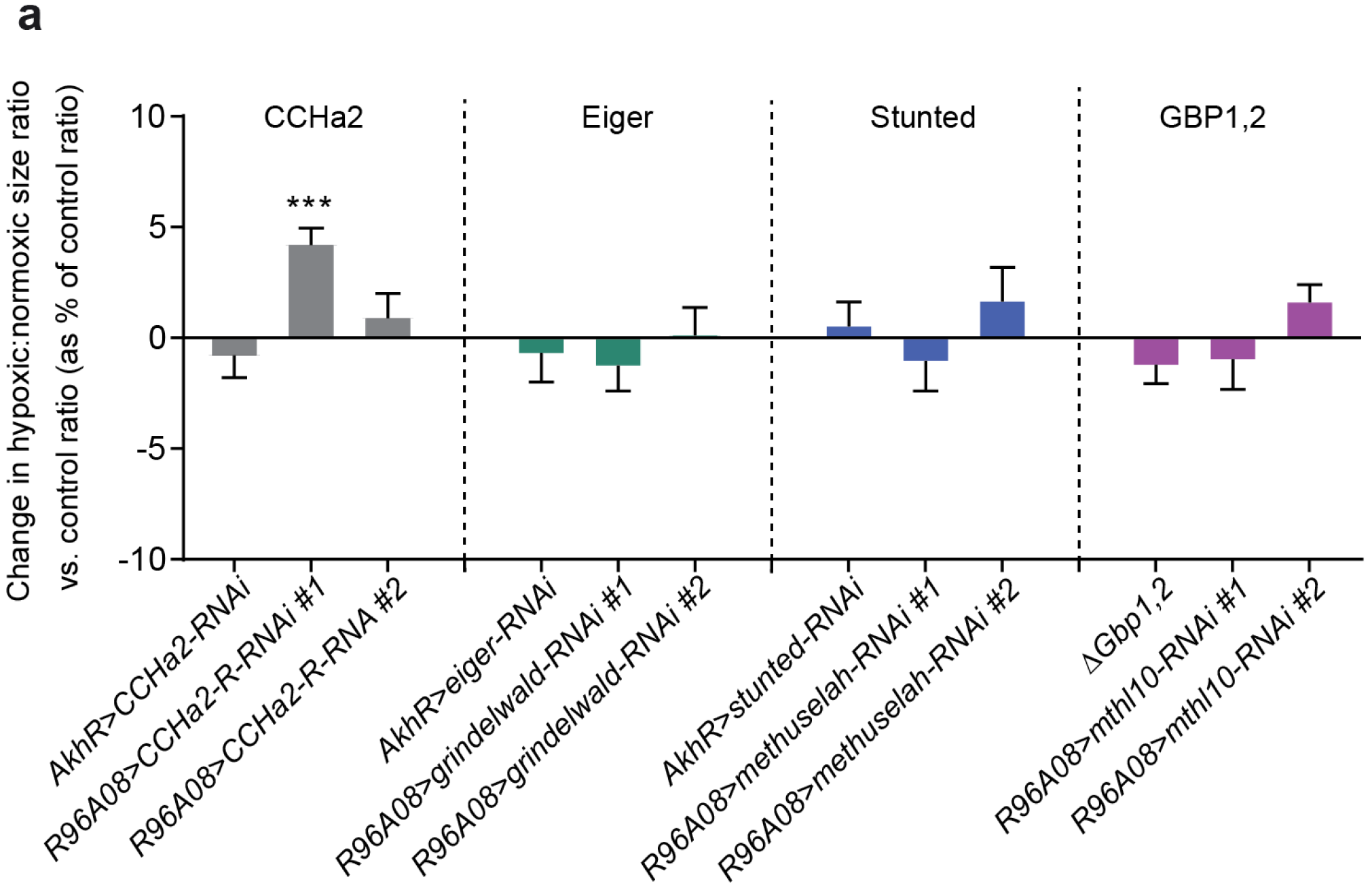
Mini-screen of fat-body factors and receptors. To assess the contributions of known fat-body factors and IPC-expressed receptors to hypoxia-induced size reduction, we assayed RNAi lines and mutants for this effect, hypothesizing that knocking down a factor or receptor that is involved in the insulinostatic response would rescue the size reduction (*i.e.*, lead to larger pupae) under hypoxia. *AkhR-GAL4∷p65* (fat body) and *R96A08-GAL4* (IPCs) drove RNAi against secreted factors and receptors, respectively. Zero change indicates that an RNAi treatment had no hypoxia-specific effect – the ratio between hypoxic and normoxic sizes was the same as for controls. Roughly +20% would represent a complete blockage of hypoxia-induced size reduction. *CCHa2/CCha2-R* results are inconsistent with each other, and a size increase upon CCHa2-R knockdown is inconsistent with its reported insulinotropic effects; thus perhaps the *CCHa2-R-RNAi-#1* phenotype reflects off-target effects. Statistics: two-tailed Student’s t-test for pairwise comparisons versus controls. ***P<0.001, compared to the *Luciferase* control.

## Acknowledgements

We thank our Erasmus-program visiting undergraduate Niyati Jain for performing the RNAi mini-screen (Fig. S6). We thank Pierre Léopold, Jan Veenstra, and Dick Nässel for DILP2, −3 and −5 antibodies, Takashi Koyama for Growth-Blocking-Peptide-related lines, Pablo Wappner for the *Hph* mutant, and Stefan Luschnig for the *GFP∷ODD* line. This work was supported by the Novo Nordisk Foundation STAR grant and an Innovation foundation grant to J.L.H and K.F.R. and by Novo Nordisk Foundation grant 16OC0021270 and Danish Council for Independent Research, Natural Sciences grant 4181-00270 to K.F.R.

## Author contributions

M.J.T., A.F.J., C.F.C., D.K.S., D.F.M.M., E.T.D., S.K.P., K.A.H., and K.F.R performed the experiments and analyzed the data. M.J.T., A.F.J., C.F.C., J.L.H., K.A.H., and K.F.R designed the experiments, and M.T., K.A.H., and K.F.R wrote the manuscript.

## References

1 Andersen, D. S., Colombani, J. & Leopold, P. Coordination of organ growth: principles and outstanding questions from the world of insects. Trends Cell Biol 23, 336–344, doi:10.1016/j.tcb.2013.03.005 (2013).

2 Danielsen, E. T., Moeller, M. E. & Rewitz, K. F. Nutrient signaling and developmental timing of maturation. Curr Top Dev Biol 105, 37–67, doi:10.1016/B978-0-12-396968-2.00002-6 (2013).

3 Tennessen, J. M. & Thummel, C. S. Coordinating growth and maturation ‑ insights from Drosophila. Curr Biol 21, R750–757, doi:10.1016/j.cub.2011.06.033 (2011).

4 Callier, V. & Nijhout, H. F. Control of body size by oxygen supply reveals size-dependent and size-independent mechanisms of molting and metamorphosis. Proc Natl Acad Sci U S A 108, 14664–14669, doi:10.1073/pnas.1106556108 (2011).

5 Callier, V. et al. The role of reduced oxygen in the developmental physiology of growth and metamorphosis initiation in Drosophila melanogaster. J Exp Biol 216, 4334–4340, doi:10.1242/jeb.093120 (2013).

6 Hietakangas, V. & Cohen, S. M. Regulation of tissue growth through nutrient sensing. Annu Rev Genet 43, 389–410, doi:10.1146/annurev-genet-102108-134815 (2009).

7 Boulan, L., Milan, M. & Leopold, P. The Systemic Control of Growth. Cold Spring Harb Perspect Biol 7, doi:10.1101/cshperspect.a019117 (2015).

8 Brogiolo, W. et al. An evolutionarily conserved function of the Drosophila insulin receptor and insulin-like peptides in growth control. Curr Biol 11, 213–221 (2001).

9 Wang, S., Tulina, N., Carlin, D. L. & Rulifson, E. J. The origin of islet-like cells in Drosophila identifies parallels to the vertebrate endocrine axis. Proc Natl Acad Sci U S A 104, 19873–19878, doi:10.1073/pnas.0707465104 (2007).

10 Ikeya, T., Galic, M., Belawat, P., Nairz, K. & Hafen, E. Nutrient-dependent expression of insulin-like peptides from neuroendocrine cells in the CNS contributes to growth regulation in Drosophila. Curr Biol 12, 1293–1300 (2002).

11 Rajan, A. & Perrimon, N. Drosophila cytokine unpaired 2 regulates physiological homeostasis by remotely controlling insulin secretion. Cell 151, 123–137, doi:10.1016/j.cell.2012.08.019 (2012).

12 Sano, H. et al. The Nutrient-Responsive Hormone CCHamide-2 Controls Growth by Regulating Insulin-like Peptides in the Brain of Drosophila melanogaster. PLoS Genet 11, e1005209, doi:10.1371/journal.pgen.1005209 (2015).

13 Sun, J. et al. Drosophila FIT is a protein-specific satiety hormone essential for feeding control. Nat Commun 8, 14161, doi:10.1038/ncomms14161 (2017).

14 Koyama, T. & Mirth, C. K. Growth-Blocking Peptides As Nutrition-Sensitive Signals for Insulin Secretion and Body Size Regulation. PLoS Biol 14, e1002392, doi:10.1371/journal.pbio.1002392 (2016).

15 Delanoue, R. et al. Drosophila insulin release is triggered by adipose Stunted ligand to brain Methuselah receptor. Science 353, 1553–1556, doi:10.1126/science.aaf8430 (2016).

16 Agrawal, N. et al. The Drosophila TNF Eiger Is an Adipokine that Acts on Insulin-Producing Cells to Mediate Nutrient Response. Cell Metab 23, 675–684, doi:10.1016/j.cmet.2016.03.003 (2016).

17 Colombani, J. et al. Antagonistic actions of ecdysone and insulins determine final size in Drosophila. Science 310, 667–670, doi:10.1126/science.1119432 (2005).

18 McBrayer, Z. et al. Prothoracicotropic hormone regulates developmental timing and body size in Drosophila. Dev Cell 13, 857–871, doi:10.1016/j.devcel.2007.11.003 (2007).

19 Mirth, C., Truman, J. W. & Riddiford, L. M. The role of the prothoracic gland in determining critical weight for metamorphosis in Drosophila melanogaster. Curr Biol 15, 1796–1807, doi:10.1016/j.cub.2005.09.017 (2005).

20 Palos, L. A. & Blasko, G. Effect of hypoxia on the development of Drosophila melanogaster (Meigen). Aviat Space Environ Med 50, 411–412 (1979).

21 Callier, V. & Nijhout, H. F. Body size determination in insects: a review and synthesis of size-and brain-dependent and independent mechanisms. Biol Rev Camb Philos Soc 88, 944–954, doi:10.1111/brv.12033 (2013).

22 Henry, J. R. & Harrison, J. F. Plastic and evolved responses of larval tracheae and mass to varying atmospheric oxygen content in Drosophila melanogaster. J Exp Biol 207, 3559–3567, doi:10.1242/jeb.01189 (2004).

23 Peck, L. S. & Maddrell, S. H. P. Limitation of size by hypoxia in the fruit flyDrosophila melanogaster. Journal of Experimental Zoology Part A: Comparative Experimental Biology 303A, 968–975, doi:10.1002/jez.a.211 (2005).

24 Frazier, M. R., Woods, H. A. & Harrison, J. F. Interactive effects of rearing temperature and oxygen on the development of Drosophila melanogaster. Physiol Biochem Zool 74, 641–650, doi:10.1086/322172 (2001).

25 Frisancho, A. R. Developmental functional adaptation to high altitude: review. Am J Hum Biol 25, 151–168, doi:10.1002/ajhb.22367 (2013).

26 Frisancho, A. R. & Baker, P. T. Altitude and growth: a study of the patterns of physical growth of a high altitude Peruvian Quechua population. Am J Phys Anthropol 32, 279–292, doi:10.1002/ajpa.1330320217 (1970).

27 Van Voorhies, W. A. Metabolic function in Drosophila melanogaster in response to hypoxia and pure oxygen. J Exp Biol 212, 3132–3141, doi:10.1242/jeb.031179 (2009).

28 Harrison, J. F., Shingleton, A. W. & Callier, V. Stunted by Developing in Hypoxia: Linking Comparative and Model Organism Studies. Physiol Biochem Zool 88, 455–470, doi:10.1086/682216 (2015).

29 Gorr, T. A., Gassmann, M. & Wappner, P. Sensing and responding to hypoxia via HIF in model invertebrates. J Insect Physiol 52, 349–364, doi:10.1016/j.jinsphys.2006.01.002 (2006).

30 Glazer, L. & Shilo, B. Z. The Drosophila FGF-R homolog is expressed in the embryonic tracheal system and appears to be required for directed tracheal cell extension. Genes Dev 5, 697–705, doi:10.1101/gad.5.4.697 (1991).

31 Klambt, C., Glazer, L. & Shilo, B. Z. breathless, a Drosophila FGF receptor homolog, is essential for migration of tracheal and specific midline glial cells. Genes Dev 6, 1668–1678, doi:10.1101/gad.6.9.1668 (1992).

32 Sutherland, D., Samakovlis, C. & Krasnow, M. A. branchless encodes a Drosophila FGF homolog that controls tracheal cell migration and the pattern of branching. Cell 87, 1091–1101, doi:10.1016/S0092-8674(00)81803-6 (1996).

33 Jarecki, J., Johnson, E. & Krasnow, M. A. Oxygen regulation of airway branching in Drosophila is mediated by branchless FGF. Cell 99, 211–220, doi:10.1016/S0092-8674(00)81652-9 (1999).

34 Centanin, L. et al. Cell autonomy of HIF effects in Drosophila: tracheal cells sense hypoxia and induce terminal branch sprouting. Dev Cell 14, 547–558, doi:10.1016/j.devcel.2008.01.020 (2008).

35 Dietzl, G. et al. A genome-wide transgenic RNAi library for conditional gene inactivation in Drosophila. Nature 448, 151–156, doi:10.1038/nature05954 (2007).

36 Ni, J. Q. et al. A genome-scale shRNA resource for transgenic RNAi in Drosophila. Nat Methods 8, 405–407, doi:10.1038/nmeth.1592 (2011).

37 Chen, C., Jack, J. & Garofalo, R. S. The Drosophila insulin receptor is required for normal growth. Endocrinology 137, 846–856, doi:10.1210/endo.137.3.8603594 (1996).

38 Loren, C. E. et al. Identification and characterization of DAlk: a novel Drosophila melanogaster RTK which drives ERK activation in vivo. Genes Cells 6, 531–544, doi:10.1046/j.1365-2443.2001.00440.x (2001).

39 Okamoto, N. & Nishimura, T. Signaling from Glia and Cholinergic Neurons Controls Nutrient-Dependent Production of an Insulin-like Peptide for Drosophila Body Growth. Dev Cell 35, 295–310, doi:10.1016/j.devcel.2015.10.003 (2015).

40 Diaz-Benjumea, F. J. & Garcia-Bellido, A. Behaviour of cells mutant for an EGF receptor homologue of Drosophila in genetic mosaics. Proc Biol Sci 242, 36–44, doi:10.1098/rspb.1990.0100 (1990).

41 Rewitz, K. F., Yamanaka, N., Gilbert, L. I. & O’Connor, M. B. The insect neuropeptide PTTH activates receptor tyrosine kinase torso to initiate metamorphosis. Science 326, 1403–1405, doi:10.1126/science.1176450 (2009).

42 Danielsen, E. T. et al. Transcriptional control of steroid biosynthesis genes in the Drosophila prothoracic gland by ventral veins lacking and knirps. PLoS Genet 10, e1004343, doi:10.1371/journal.pgen.1004343 (2014).

43 Bialecki, M., Shilton, A., Fichtenberg, C., Segraves, W. A. & Thummel, C. S. Loss of the ecdysteroid-inducible E75A orphan nuclear receptor uncouples molting from metamorphosis in Drosophila. Dev Cell 3, 209–220, doi:S1534580702002046 [pii] (2002).

44 Huang, X., Warren, J. T., Buchanan, J., Gilbert, L. I. & Scott, M. P. Drosophila Niemann-Pick type C-2 genes control sterol homeostasis and steroid biosynthesis: a model of human neurodegenerative disease. Development 134, 3733–3742, doi:10.1242/dev.004572 (2007).

45 Wilk, R., Weizman, I. & Shilo, B. Z. trachealess encodes a bHLH-PAS protein that is an inducer of tracheal cell fates in Drosophila. Genes Dev 10, 93–102, doi:10.1101/gad.10.1.93 (1996).

46 Ohshiro, T., Emori, Y. & Saigo, K. Ligand-dependent activation of breathless FGF receptor gene in Drosophila developing trachea. Mech Dev 114, 3–11, doi:10.1016/S0925-4773(02)00042-4 (2002).

47 Britton, J. S., Lockwood, W. K., Li, L., Cohen, S. M. & Edgar, B. A. Drosophila’s insulin/PI3-kinase pathway coordinates cellular metabolism with nutritional conditions. Dev Cell 2, 239–249, doi:10.1016/S1534-5807(02)00117-X (2002).

48 Kim, J. & Neufeld, T. P. Dietary sugar promotes systemic TOR activation in Drosophila through AKH-dependent selective secretion of Dilp3. Nat Commun 6, 6846, doi:10.1038/ncomms7846 (2015).

49 Puig, O. & Tjian, R. Transcriptional feedback control of insulin receptor by dFOXO/FOXO1. Genes Dev 19, 2435–2446, doi:10.1101/gad.1340505 (2005).

50 Mirth, C. K. & Riddiford, L. M. Size assessment and growth control: how adult size is determined in insects. Bioessays 29, 344–355, doi:10.1002/bies.20552 (2007).

51 Jones, T. A. & Metzstein, M. M. Examination of Drosophila larval tracheal terminal cells by light microscopy. J Vis Exp, e50496, doi:10.3791/50496 (2013).

52 Nambu, J. R., Chen, W., Hu, S. & Crews, S. T. The Drosophila melanogaster similar bHLH-PAS gene encodes a protein related to human hypoxia-inducible factor 1 alpha and Drosophila single-minded. Gene 172, 249–254, doi:10.1016/0378-1119(96)00060-1 (1996).

53 Bacon, N. C. et al. Regulation of the Drosophila bHLH-PAS protein Sima by hypoxia: functional evidence for homology with mammalian HIF-1 alpha. Biochem Biophys Res Commun 249, 811–816, doi:10.1006/bbrc.1998.9234 (1998).

54 Centanin, L., Ratcliffe, P. J. & Wappner, P. Reversion of lethality and growth defects in Fatiga oxygen-sensor mutant flies by loss of hypoxia-inducible factor-alpha/Sima. EMBO Rep 6, 1070–1075, doi:10.1038/sj.embor.7400528 (2005).

55 Frei, C. & Edgar, B. A. Drosophila cyclin D/Cdk4 requires Hif-1 prolyl hydroxylase to drive cell growth. Dev Cell 6, 241–251, doi:10.1016/S1534-5807(03)00409-X (2004).

56 Bruick, R. K. & McKnight, S. L. A conserved family of prolyl-4-hydroxylases that modify HIF. Science 294, 1337–1340, doi:10.1126/science.1066373 (2001).

57 Geminard, C., Rulifson, E. J. & Leopold, P. Remote control of insulin secretion by fat cells in Drosophila. Cell Metab 10, 199–207, doi:10.1016/j.cmet.2009.08.002 (2009).

58 Misra, T. et al. A genetically encoded biosensor for visualising hypoxia responses in vivo. Biol Open 6, 296–304, doi:10.1242/bio.018226 (2017).

59 Wong, D. M., Shen, Z., Owyang, K. E. & Martinez-Agosto, J. A. Insulin-and warts-dependent regulation of tracheal plasticity modulates systemic larval growth during hypoxia in Drosophila melanogaster. PLoS One 9, e115297, doi:10.1371/journal.pone.0115297 (2014).

60 Krock, B. L., Skuli, N. & Simon, M. C. Hypoxia-induced angiogenesis: good and evil. Genes Cancer 2, 1117–1133, doi:10.1177/1947601911423654 (2011).

61 Fong, G. H. Mechanisms of adaptive angiogenesis to tissue hypoxia. Angiogenesis 11, 121–140, doi:10.1007/s10456-008-9107-3 (2008).

62 Colombani, J. et al. A nutrient sensor mechanism controls Drosophila growth. Cell 114, 739–749, doi:10.1016/S0092-8674(03)00713-X (2003).

63 Gronke, S., Clarke, D. F., Broughton, S., Andrews, T. D. & Partridge, L. Molecular evolution and functional characterization of Drosophila insulin-like peptides. PLoS Genet 6, e1000857, doi:10.1371/journal.pgen.1000857 (2010).

64 Xu, G. G. & Rothenberg, P. L. Insulin receptor signaling in the beta-cell influences insulin gene expression and insulin content: evidence for autocrine beta-cell regulation. Diabetes 47, 1243–1252, doi:10.2337/diab.47.8.1243 (1998).

65 Smith, S. R. & Ravussin, E. Role of Adipocyte in Metabolism and Endocrine Function. (Elsevier Saunders, 2006).

66 Regazzetti, C. et al. Hypoxia decreases insulin signaling pathways in adipocytes. Diabetes 58, 95–103, doi:10.2337/db08-0457 (2009).

67 Pasarica, M. et al. Reduced adipose tissue oxygenation in human obesity: evidence for rarefaction, macrophage chemotaxis, and inflammation without an angiogenic response. Diabetes 58, 718–725, doi:10.2337/db08-1098 (2009).

68 Polotsky, V. Y. et al. Intermittent hypoxia increases insulin resistance in genetically obese mice. J Physiol 552, 253–264, doi:10.1113/jphysiol.2003.048173 (2003).

69 Norouzirad, R., Gonzalez-Muniesa, P. & Ghasemi, A. Hypoxia in Obesity and Diabetes: Potential Therapeutic Effects of Hyperoxia and Nitrate. Oxid Med Cell Longev 2017, 5350267, doi:10.1155/2017/5350267 (2017).

70 Rasouli, N. Adipose tissue hypoxia and insulin resistance. J Investig Med 64, 830–832, doi:10.1136/jim-2016-000106 (2016).

71 Hosogai, N. et al. Adipose tissue hypoxia in obesity and its impact on adipocytokine dysregulation. Diabetes 56, 901–911, doi:10.2337/db06-0911 (2007).

72 Lefere, S. et al. Hypoxia-regulated mechanisms in the pathogenesis of obesity and non-alcoholic fatty liver disease. Cell Mol Life Sci 73, 3419–3431, doi:10.1007/s00018-016-2222-1 (2016).

73 Kihira, Y. et al. Deletion of hypoxia-inducible factor-1alpha in adipocytes enhances glucagon-like peptide-1 secretion and reduces adipose tissue inflammation. PLoS One 9, e93856, doi:10.1371/journal.pone.0093856 (2014).

74 Ryan, S. Adipose tissue inflammation by intermittent hypoxia: mechanistic link between obstructive sleep apnoea and metabolic dysfunction. J Physiol 595, 2423–2430, doi:10.1113/JP273312 (2017).

75 Masson, N. & Ratcliffe, P. J. Hypoxia signaling pathways in cancer metabolism: the importance of co-selecting interconnected physiological pathways. Cancer Metab 2, 3, doi:10.1186/2049-3002-2-3 (2014).

76 Reiling, J. H. & Hafen, E. The hypoxia-induced paralogs Scylla and Charybdis inhibit growth by down-regulating S6K activity upstream of TSC in Drosophila. Genes Dev 18, 2879–2892, doi:10.1101/gad.322704 (2004).

77 Brugarolas, J. et al. Regulation of mTOR function in response to hypoxia by REDD1 and the TSC1/TSC2 tumor suppressor complex. Genes Dev 18, 2893–2904, doi:10.1101/gad.1256804 (2004).

78 Cantley, J. The control of insulin secretion by adipokines: current evidence for adipocyte-beta cell endocrine signalling in metabolic homeostasis. Mamm Genome 25, 442–454, doi:10.1007/s00335-014-9538-7 (2014).

79 Wu, L. E. et al. Identification of fatty acid binding protein 4 as an adipokine that regulates insulin secretion during obesity. Mol Metab 3, 465–473, doi:10.1016/j.molmet.2014.02.005 (2014).

80 Pfeiffer, B. D. et al. Refinement of tools for targeted gene expression in Drosophila. Genetics 186, 735–755, doi:10.1534/genetics.110.119917 (2010).

81 Venken, K. J. et al. Versatile P[acman] BAC libraries for transgenesis studies in Drosophila melanogaster. Nat Methods 6, 431–434, doi:10.1038/nmeth.1331 (2009).

82 Lin, D. M. & Goodman, C. S. Ectopic and increased expression of Fasciclin II alters motoneuron growth cone guidance. Neuron 13, 507–523, doi:10.1016/0896-6273(94)90022-1 (1994).

83 Ranganayakulu, G., Schulz, R. A. & Olson, E. N. Wingless signaling induces nautilus expression in the ventral mesoderm of the Drosophila embryo. Dev Biol 176, 143–148, doi:10.1006/dbio.1996.9987 (1996).

84 Pfeiffer, B. D. et al. Tools for neuroanatomy and neurogenetics in Drosophila. Proc Natl Acad Sci U S A 105, 9715–9720, doi:10.1073/pnas.0803697105 (2008).

85 Tapon, N., Ito, N., Dickson, B. J., Treisman, J. E. & Hariharan, I. K. The Drosophila tuberous sclerosis complex gene homologs restrict cell growth and cell proliferation. Cell 105, 345–355, doi:10.1016/S0092-8674(01)00332-4 (2001).

86 Honegger, B. et al. Imp-L2, a putative homolog of vertebrate IGF-binding protein 7, counteracts insulin signaling in Drosophila and is essential for starvation resistance. J Biol 7, 10, doi:10.1186/jbiol72 (2008).

87 Hu, Y., Comjean, A., Perkins, L. A., Perrimon, N. & Mohr, S. E. GLAD: an Online Database of Gene List Annotation for Drosophila. J Genomics 3, 75–81, doi:10.7150/jgen.12863 (2015).

88 Zerbino, D. R. et al. Ensembl 2018. Nucleic Acids Res 46, D754–D761, doi:10.1093/nar/gkx1098 (2018).

89 Moeller, M. E. et al. Warts Signaling Controls Organ and Body Growth through Regulation of Ecdysone. Curr Biol 27, 1652–1659 e1654, doi:10.1016/j.cub.2017.04.048 (2017).

90 Schindelin, J. et al. Fiji: an open-source platform for biological-image analysis. Nature methods 9, 676–682, doi:10.1038/nmeth.2019 (2012).

91 Soderberg, J. A., Birse, R. T. & Nassel, D. R. Insulin production and signaling in renal tubules of Drosophila is under control of tachykinin-related peptide and regulates stress resistance. PLoS One 6, e19866, doi:10.1371/journal.pone.0019866 (2011).

92 Danielsen, E. T. et al. A Drosophila Genome-Wide Screen Identifies Regulators of Steroid Hormone Production and Developmental Timing. Dev Cell 37, 558–570, doi:10.1016/j.devcel.2016.05.015 (2016).

93 Hentze, J. L., Carlson, M. A., Nassel, D. R. & Rewitz, K. F. The neuropeptide Allatostatin A coordinates feeding decisions and metabolism in Drosophila. Submitte to Plos Genetics (2013).

